# Distribution theories for genetic line of least resistance and evolvability measures

**DOI:** 10.1101/2023.10.07.561348

**Authors:** Junya Watanabe

## Abstract

Quantitative genetic theory on multivariate character evolution predicts that a population’s response to directional selection tends to happen around the major axis of the genetic covariance matrix **G**—the so-called genetic line of least resistance. Inferences on the genetic constraints in this sense have traditionally been made by measuring the angle of deviation of evolutionary trajectories from the major axis, or more recently by calculating the amount of genetic variance—the Hansen–Houle evolvability—available along the trajectories. However, there have not been clear practical guidelines on how these quantities can be interpreted, especially in a high-dimensional space. This study summarizes pertinent distribution theories for relevant quantities, pointing out that they can be written as ratios of quadratic forms in evolutionary trajectory vectors by taking **G** as a parameter. For example, a beta distribution with appropriate parameters can be used as a null distribution for squared cosine of the angle of deviation from a major axis or subspace. More general cases can be handled with the probability distribution of ratios of quadratic forms in normal variables. Apart from its use in hypothesis-testing, this latter approach could potentially be used as a heuristic tool for looking into various selection scenarios like directional and/or correlated selection as parameterized with mean and covariance of selection gradients.

## 1 Introduction

The predictability of phenotypic evolution has long been a central topic in evolutionary biology. The quantitative genetic theory of multivariate continuous character evolution provides a particularly useful predictive framework in the sense that theoretical prediction is amenable to empirical test. The theory states that, when a population is subject to directional selection on *p*-dimensional traits under certain simplifying conditions, its response can be described by the Lande equation (Lande, 1979; Lande & Arnold, 1983):

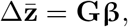

where 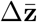 is a *p*-dimensional vector of across-generational change in the mean phenotype, **G** is the *p ×p* additive genetic covariance matrix of the traits, and **β** is a *p*-dimensional selection gradient vector, which is empirically measured as partial regression coefficients of trait values to the relative fitness. A consequence of this relationship is that the response 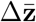 is in general not perfectly aligned with the direction of selection **β**, but tends to be more biased or constrained toward the major axis of character covariation entailed in **G**. This theory has been playing a pivotal role in the current research on quantitative character evolution (Arnold *et al*., 2001; Steppan *et al*., 2002; Walsh & Blows, 2009; Hansen & Pélabon, 2021).

The above theory as originally formulated pertained to microevolutionary changes across successive generations. It was the seminal paper of Schluter (1996) that greatly extended the scope into the predictability of long-term multivariate phenotypic evolution using this theory. He conjectured that, if selection happens toward an optimal state that is stable across multiple generations and phenotypic evolution proceeds as the Lande equation predicts, evolutionary change in a multivariate trait space should be concentrated in the direction of the major axis of genetic (co)variation between the traits, especially in an early stage of adaptive evolution. If the model holds, the rate of phenotypic evolution should also be faster near the direction of the major axis. Empirical tests of these hypotheses involve measuring the major axis of genetic variation, called the genetic line of least resistance **g**_max_, and the angle of deviation *θ* of evolutionary trajectory or divergence vectors from it (Fig. 1). The former is represented by the eigenvector of **G** corresponding to the largest eigenvalue, which tends to (although not necessarily) explain a predominantly large proportion of total variance. Following this conceptualization, extensive investigations have been conducted to test the role of genetic constraints in long-term evolution, in the sense that empirical phenotypic divergences between populations or species are aligned with (proxies of) **g**_max_ in ancestors (e.g., Bégin & Roff, 2003, 2004; Marroig & Cheverud, 2005; McGuigan *et al*., 2005; Renaud *et al*., 2006; Berner *et al*., 2010; Renaud & Auffray, 2013; McGlothlin *et al*., 2018; Royauté *et al*., 2020; Henry & Stinchcombe, 2023a).

**Figure 1.**
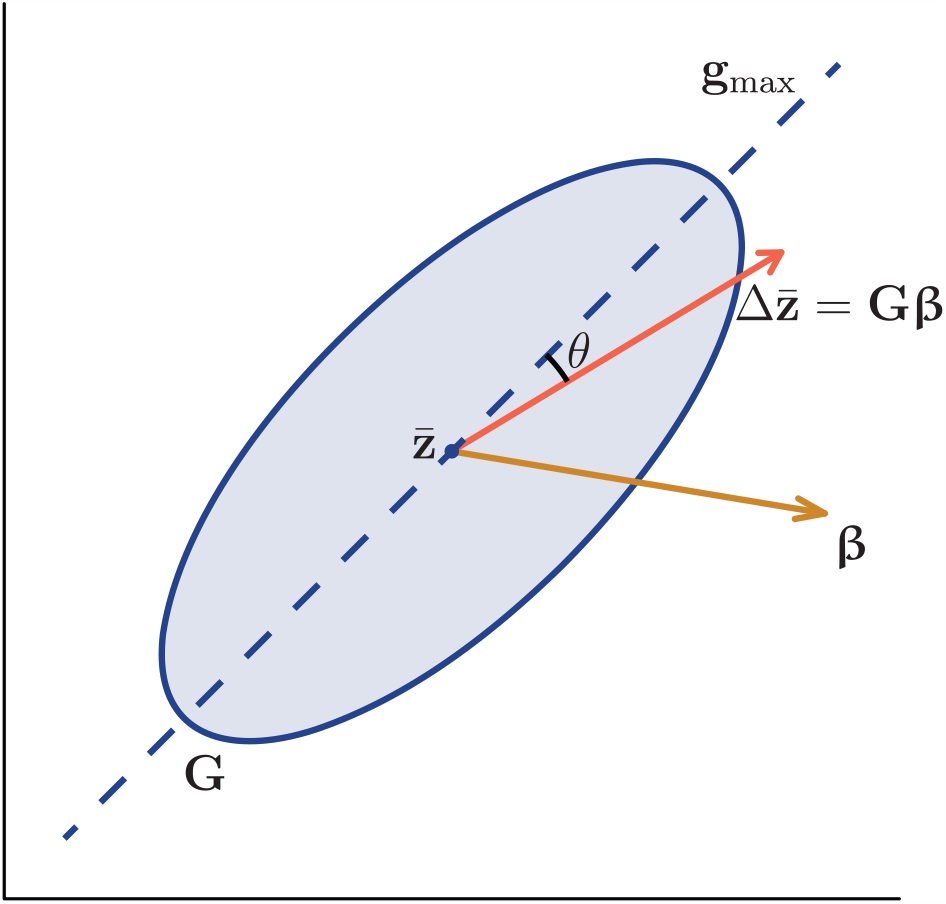
Genetic line of least resistance **g**_max_ and angle of deviation *θ*. Here, the genetic covariance matrix **G** is depicted by an ellipse of equiprobability contour, selection gradient **β** by a gold narrow-headed arrow, and response 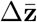 by an orange broad-headed arrow. **g**_max_ is the eigenvector of **G** corresponding to the largest eigenvalue, from which the angle of deviation *θ* of 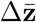 is typically measured.

More recently, the use of the angle of deviation from **g**_max_ for testing a constraint hypothesis has been criticized by subsequent authors (Blows & Higgie, 2003; Hansen & Voje, 2011), who pointed out that there can in general be more dimensions with large genetic variance than can be represented by a single eigenvector. As an alternative, a few studies (Hunt, 2007; Grabowski *et al*., 2011; Hansen & Voje, 2011) independently proposed to look at the amount of genetic variance available in the direction of evolutionary trajectories—evolvability of Hansen & Houle (2008). This alternative measure is arguably more biologically meaningful than deviation from **g**_max_, and has been utilized in many subsequent studies (e.g., Haber, 2016; Baab, 2018; Opedal *et al*., 2022).

Together, these lines of studies can be seen as a trend to connect the phenotypic divergence patterns at the inter-population level to the phenotypic variability at the intra-population level. There have been criticisms and caveats from various standpoints over this research program, including its blindness to mechanistic basis, simplistic assumptions, and uncertainty in the stability of **G** over pertinent timescales (Pigliucci, 2006; Morrissey *et al*., 2010; Milocco & Salazar-Ciudad, 2020, 2022; Henry & Stinchcombe, 2023b). Nevertheless, active research effort exemplifies persistent interest in understanding potential roles of constraints in macroevolutionary phenotypic diversification (Polly & Mock, 2018; Watanabe, 2018; Machado, 2020; Renaud *et al*., 2021; Rhoda *et al*., 2023). In addition, recent meta-analyses have demonstrated that the amount of evolvability available to selection can roughly predict that of phenotypic divergence (Opedal *et al*., 2023; Voje *et al*., 2023).

This methodological paper focuses a pragmatic aspect of this research program, that is, the probability distributions of relevant statistics. Inferences and tests on genetic constraint hypotheses using the angle of deviation or evolvability have often been conducted by comparing, formally or informally, observed values with their hypothetical distributions. Empirical procedures varied from such naïve ones like face-value interpretation of angles, where large deviations were interpreted as evidence for a limited role of constraints (e.g., Eroukhmanoff & Svensson, 2011), to more or less sophisticated ones like comparison with the theoretical range and mean (Hansen & Voje, 2011) or with Monte Carlo distributions (Renaud *et al*., 2006; Hunt, 2007). These procedures are incomplete in the sense that they do not take into account the theoretical probability distributions of the focal statistics, theories on which have indeed been lacking altogether in the evolutionary quantitative genetics literature. This paper supplements this lack of knowledge by gathering useful results from the statistical literature. In particular, it makes use of the fact that focal statistics—squared cosine and projected variance—can be written as a ratio of quadratic forms, whose distribution can be fully characterized under certain assumptions. The theories provide a guide for interpretation of, and an accurate means for statistical hypothesis testing on, the focal quantities.

## 2 Theory

### 2.1 Preliminaries

To start with, basic definitions of the angle in a vector space is reviewed. Consider the two non-zero vectors **x** = (*x*_1_, …, *x*_*p*_)^*T*^ and **y** = (*y*_1_, …, *y*_*p*_)^*T*^, where the superscript ^*T*^ denotes matrix transpose, in a *p*-dimensional Euclidean space. The angle *θ* between these two vectors can be defined from

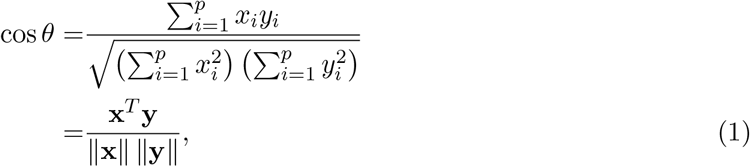

where ∥·∥ denotes the vector norm or length: 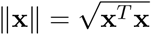. In other words, *θ* is the arc cosine of the inner product of the two vectors standardized by their norms. Note that cos *θ* is also the Pearson product-moment correlation coefficient between **x** and **y** without mean-centering, which is known as the vector correlation in the biological literature. The Cauchy–Schwarz inequality dictates that the right-hand side of (1) is bounded within the interval [*−*1, 1]. Following the convention, the range of *θ* is taken as [0, *π*]; *θ* = 0 (cos *θ* = 1) corresponds to the positive proportionality or complete alignment of the two vectors, *θ* = π */*2 (cos *θ* = 0) to the orthogonality, and *θ* = *π* (cos *θ* = *−*1) to the negative proportionality. An assumption here is that both **x** and **y** have fixed polarities.

For the present purpose, however, one concerns the angle between a random vector and an axis, or more generally a subspace, spanned by eigenvectors, whose polarities are arbitrary by definition. As such, it is more natural to consider the squared cosine of the angle.^1^ For generality, consider the angle *θ* between a *p*-variate random vector **x** and the *q*-dimensional subspace spanned by the columns of a *p ×q* full-column-rank matrix **V** (Fig. 2). Hereafter, this subspace is denoted by the range *R* (**V**) = *{***Vx** : **x** *∈* R^q^ *}*. In most typical applications, the columns of **V** are the eigenvectors of **G** corresponding to the *q* largest eigenvalues (see, e.g., Blows *et al*., 2004) (i.e., **V** = **g**_max_ when *q* = 1), but this setting can accommodate any subspace of interest that is even not necessarily bound to eigenvectors of **G**. Here, cos^2^ *θ* is the ratio of the squared norm of the vector made by orthogonally projecting **x** onto the subspace to that of the original vector **x** itself (Fig. 2). Algebraically, the orthogonal projection to this subspace is represented by (e.g., Schott, 2016, chapter 2)

**Figure 2.**
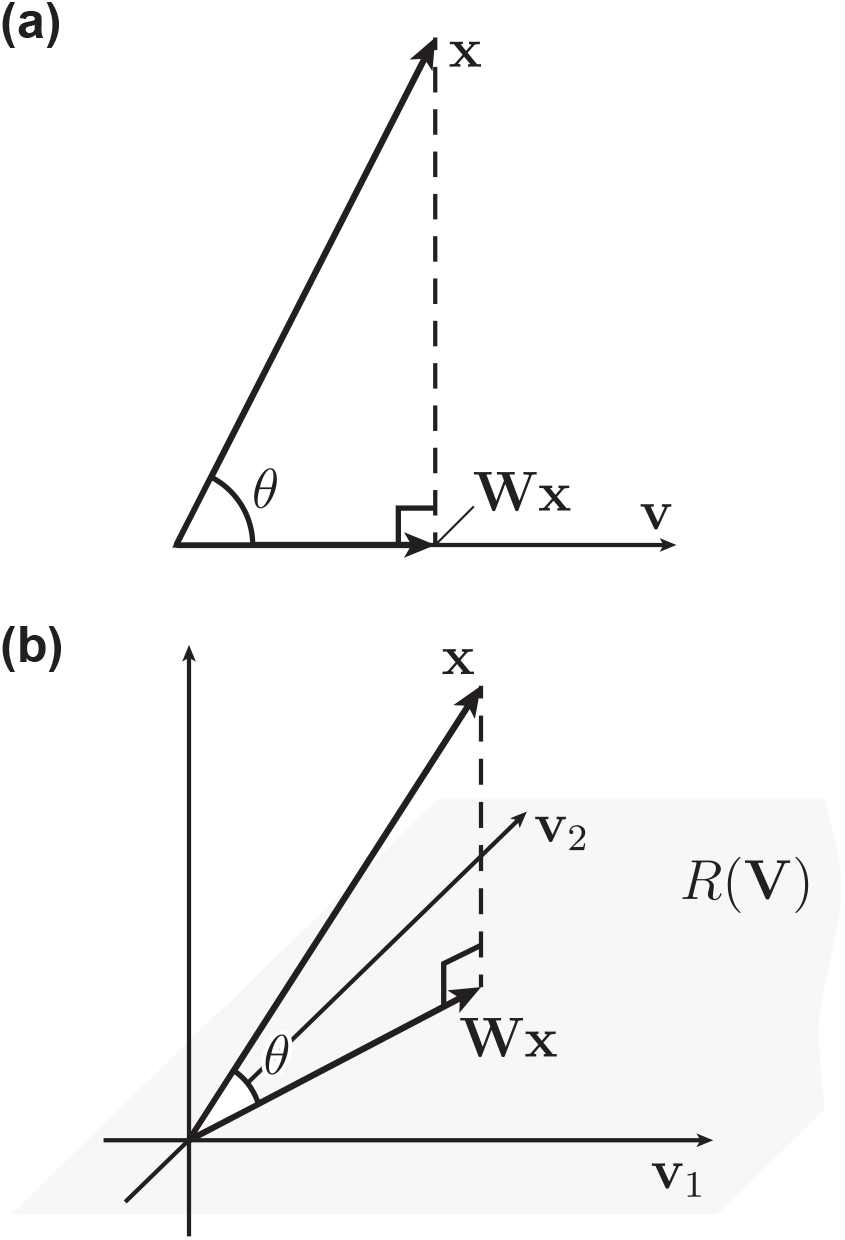
Angle of deviation *θ* of a random vector **x** from the subspace spanned by (**a**) a single vector **v** (*q* = 1) or (**b**) two vectors **v**_1_ and **v**_2_ (*q* = 2). In **b**, the subspace is denoted by the range *R*(**V**), with **V** being a matrix that has the two vectors as columns. The projection matrix **W** formed from **V** (2) projects any vector in the space orthogonally onto *R*(**V**).

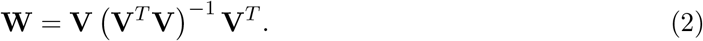

When the columns of **V** are orthogonal (as is the case for eigenvectors of symmetric matrices) and normalized to have unit length, **V**^*T*^ **V** = **I**_*q*_, the *q*-dimensional identity matrix, so that the expression simplifies into **W** = **VV**^*T*^ . To demystify, assume the simplest situation that the columns of **V** correspond to the first *q* coordinate axes. In that case, **W** = diag (1, …, 1, 0, …, 0) with *q* ones and *p − q* zeros along the diagonal, so it just extracts the first *q* components from a vector, replacing the other components with zeros. These ones and zeros are the only possible combination of eigenvalues for **W**, regardless of the choice of **V** given *p, q* (see also Schott, 2016, chapter 11). With this projection matrix, and noting **W**^2^ = **W**,

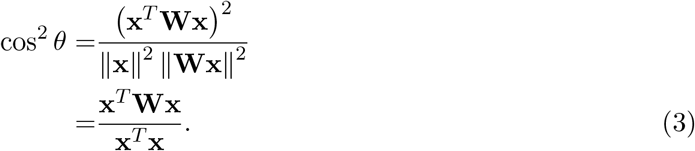

Thus, the squared cosine is a ratio of quadratic forms in **x**, for which lots of useful results are available as seen below. Hereafter, this quantity is denoted by 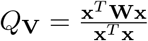 for brevity.

The projection matrix **W** has often been used in the biological literature (e.g., Burnaby, 1966; Blows *et al*., 2004). In the terminology of linear regression, **W** is also known as the hat matrix, which transforms the vector of response variable to that of predicted values. Also, *Q*_**V**_ can be called a squared correlation (*q* = 1) or squared multiple correlation (*q >* 1) coefficient. These are simple consequences of the fact that linear regression is geometrically an orthogonal projection of the vector of response variable to the subspace spanned by the explanatory variables in the observation space (e.g., Anderson, 2003, section 4.4).

Throughout this paper, **G** is assumed to be a validly constructed covariance matrix, i.e., to be symmetric and nonnegative definite. For the present theoretical analyses, it is further assumed to be known without error. Although the latter assumption is never satisfied in empirical analyses, most of the theoretical principles will still apply. See Discussion for details.

### 2.2 Null distribution: uniformly distributed vector

Interest is often in rejecting the null hypothesis that the alignment between an observed divergence vector and the focal subspace is no different from what is expected from the uniform distribution of the vector on the unit hypersphere in the *p*-dimensional space (hereafter *S*^*p−*1^). If the observed cosine falls above the 95 percentile, say, of a null distribution, the null hypothesis is rejected and the alternative hypothesis of alignment is accepted. In evolutionary quantitative genetics, it seems popular, if not predominant, to generate a null distribution by Monte Carlo simulation in which a large number of random vectors uniformly distributed on *S*^*p−*1^ is generated on computer (e.g., Renaud *et al*., 2006; Hunt, 2007).^2^ But a more general analytic result is readily available using the mathematical structure of *Q*_**V**_.

Note that the cosine between two vectors uniformly distributed on *S*^*p−*1^ has the same distribution as that between one uniformly distributed vector and a fixed reference vector. Technically, this equivalence is an immediate consequence of the definition of the uniform distribution on *S*^*p−*1^: invariance against orthogonal rotation (e.g., Mardia & Jupp, 1999). As constructed above, the subsequent results apply to the squared cosine between a vector uniformly distributed on *S*^*p−*1^ and any *q*-dimensional subspace of the *p*-dimensional Euclidean space. In addition, it is noted that the distribution of *Q*_**V**_ is identical under (i) the uniform distribution of **x** on *S*^*p−*1^ and (ii) the standard normal distribution of **x**: **x** *∼ N*_*p*_(**0**_*p*_, **I**_*p*_), where **0**_*p*_ is the *p*-dimensional vector of 0’s. This is because the component of magnitude *∥***x***∥* cancels between the numerator and denominator. Indeed, any spherically symmetric distribution of **x** yields the same distribution of *Q*_**V**_ in this case.

To see the distribution of squared cosine, observe

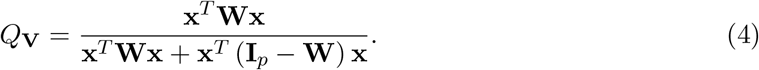

When **x** *∼ N*_*p*_(**0**_*p*_, **I**_*p*_), it is known that 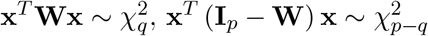, and that these are mutually independent (e.g., Schott, 2016, theorems 11.9 and 11.14). (This result is most easily seen when **W** = diag (1, …, 1, 0, …, 0), when these quantities are sums of squares of independent standard normal variables.) It is a standard result (e.g., Johnson *et al*., 1995, chapter 25) that the proportion of one chi-square variable within a sum of independent chi-square variables follows a beta distribution, with the parameters corresponding to the degrees of freedom, i.e.,

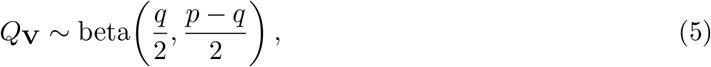

whose mean is E (*Q*_**V**_) = *q/p*. Note that this distribution does not depend on the choice of **V**, because the uniform distribution of **x** is invariant against any orthogonal rotation.

This distribution is shown in Fig. 3a for *q* = 1 and varying *p*. It is evident that the distribution of squared cosine is increasingly concentrated around 0 (or *θ* = *π /*2) as *p* increases. To repeat the popular quote, the angle between random vectors approaches the right angle as the dimensionality of the space increases.

**Figure 3.**
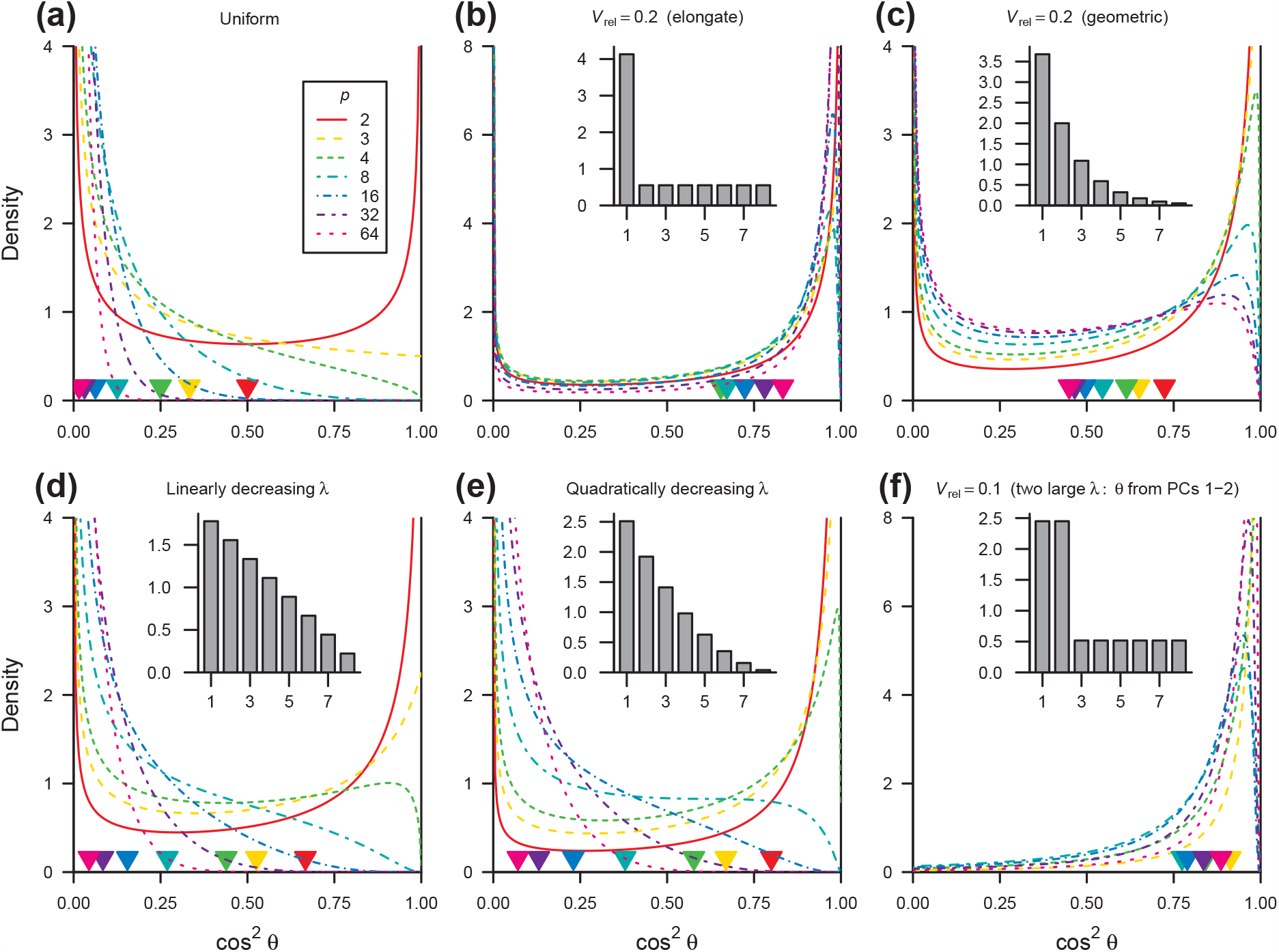
Example distributions of squared cosine. (**a**) Null distribution with a uniformly distributed vector and a fixed axis (5). (**b–e**) Distributions with a response vector and the major axis of variation (6) under the Lande model under various eigenstructures, assuming uniformly distributed selection. (**f**) Same, but deviation is measured from the plane spanned by the eigenvectors corresponding to two equally large eigenvalues. Lines represent probability densities, and triangles at the bottom represent the means, across varying dimensionality *p*. Inset scree plots represent eigenvalue conformations of **G** at *p* = 8.

This distributional result is fairly well recognized in the literature, and there are many possible ways to prove it, especially when *q* = 1 (see, e.g., Muirhead, 1982; Rice, 1990; Anderson, 2003; Cai *et al*., 2013). Watanabe (2022a) restated one of those proofs in a context where the polarities of the vectors are relevant. The result for the special case of *q* = 1 is frequently used for testing alignment between allometric vectors in the morphometric literature (e.g., Klingenberg & Marugán-Lobón, 2013), where Li (2011) seems to be a preferred citation. However, this particular result had long been known, at least dating back to Fisher (1915) (for history see Johnson *et al*., 1995).

As the distribution function of the beta distribution is available in most popular statistical packages (e.g., `stats::pbeta()` function in R; R Core Team, 2023), the oft-practiced Monte-Carlo-based test for this distribution is superfluous or even undesirable because of the randomness. For testing the null hypothesis of uniform distribution of divergence vectors, it is recommended to utilize the analytic result outlined here.

### 2.3 Distribution under Lande model

Testing the above null hypothesis may not be of interest in every situation. One may have particular biological model about the null distribution of the response vectors (Renaud & Auffray, 2013; Reddiex & Chenoweth, 2021). Or interest may be in examining whether an observed response vector conforms with the Lande model (Berner *et al*., 2010; Eroukhmanoff & Svensson, 2011). It is impossible to examine all hypotheses of potential interest, but it will be instructive to consider the distribution of squared cosine *Q*_**V**_ under the Lande model; that is, assuming **G** to remain constant and the Lande equation 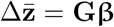 to hold across evolutionary time, with 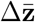 and **β** representing the net phenotypic change and selection gradient, respectively.

Inserting 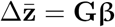 into **x** in (3) and noting the symmetry of **G**, one has

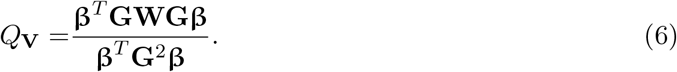

In typical applications, the columns of **V** are the eigenvectors corresponding to the *q* largest eigenvalues of **G**, in which case the expression becomes

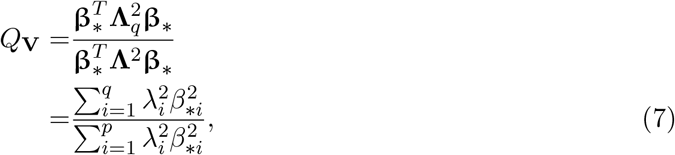

where **Λ** = diag (*λ*_1_, …, *λ*_*p*_) is a diagonal matrix of the eigenvalues of **G, Λ**_*q*_ = diag (*λ*_1_, …, *λ*_*q*_, 0, …, 0) with the *q* largest eigenvalues of **G** and *p − q* zeros, and **β***∗* = **U**^*T*^ **β** = (*β*_*∗*1_, …, *β*_*∗p*_)^*T*^ is **β** transformed by the matrix of eigenvectors **U** of **G**.

Although the probability distribution of *Q*_**V**_ in general conditions does not fall within the family of standard distributions, analytic results are available for the moments (mean, variance, etc.) and the distribution and density functions of a ratio of quadratic forms by assuming the multivariate normality of the vector: **β** *∼ N*_*p*_(**µ, Σ**) (Appendix A). This assumption is admittedly a simplistic one, but can in principle accommodate various scenarios of potential biological interest, from uniformly distributed selection (**µ** = **0**_*p*_ and **Σ** = **I**_*p*_) to directional (**µ***/ ≠* **0**_*p*_) and/or correlated (**Σ** *≠* **I**_*p*_) selection (see Chevin, 2013, for a similar parameterization). Under this assumption, the analytic results can then be used for statistical inferences, for example calculating *P* -values for individual observations or evaluating goodness of fit for a sample. Note also that 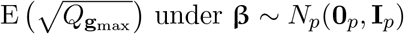 evaluated in this way provides an exact expression for mean constraints, which has been used as a descriptive measure of genetic constraints (Marroig *et al*., 2009; Melo *et al*., 2016) (Appendix A).

Example distributions of *Q*_**V**_ under the uniform distribution of selection gradients **β** *∼ N*_*p*_(**0**_*p*_, **I**_*p*_) are shown in Fig. 3b–f, where the columns of **V** are taken as the first eigenvector (Fig. 3b–e) or the first two eigenvectors (Fig. 3f) of **G**. The example eigenvalue conformations are constructed so that the relative eigenvalue variance *V*_rel_ of **G** remains constant (Fig. 3b, c, and f) or trailing eigenvalues decrease linearly or quadratically in magnitude (Fig. 3d and e) (see Watanabe, 2022b, for algorithms to construct eigenvalue conformations). As expected from the model, the response vectors are more strongly aligned with the focal subspace than in the uniform case (Fig. 3a). In the conditions where the eigenvalues are gradually decreasing in magnitude, the distributions tend to be increasingly concentrated around 0 as the dimensionality increases (Fig. 3c–e). On the other hand, in the conditions where the trailing eigenvalues are identical in magnitude, the distributions tend to be slightly more concentrated around 1 as the dimensionality increases (Fig. 3b, f). Therefore, the distribution of *Q*_**V**_ depends both on the dimensionality *p* and the eigenvalue conformations of **G**, making comparisons between different datasets rather difficult.

### 2.4 Alternative: distribution of evolvability

The use of **g**_max_ has been criticized because the axis spanned by a single eigenvector does not necessarily exhaust the dimensions with high evolvability (Blows & Higgie, 2003; Hansen & Voje, 2011). That issue can partially be mitigated by using the subspace approach described above, provided that one can decide how many axes should be incorporated (e.g., the first few eigenvectors of **G**). Nevertheless, the alternative proposed by Hunt (2007) and Hansen & Voje (2011), that is to measure the amount of genetic variance or evolvability along trajectory vectors, seems to provide a natural approach that is free from such a choice.

For this purpose, Hansen & Voje (2011) proposed to use the measures of evolvability *e* and conditional evolvability *c*, which are defined as the amounts of genetic variance available in the direction of a vector **x** in the absence and presence, respectively, of stabilizing selection in the other directions. (An equivalent of the former was called projected variance in Hunt (2007).) The expressions are (Hansen & Houle, 2008; Watanabe, 2023a)

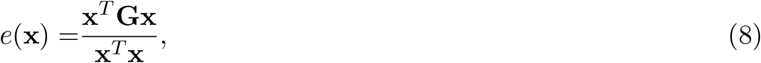

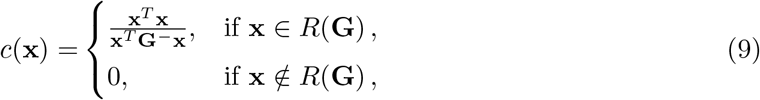

where **G**^*−*^ is a generalized inverse of **G**, which equals the ordinary inverse **G**^*−*1^ when **G** is nonsingular (see, e.g., Schott, 2016, chapter 5). The condition **x** ∉ *R*(**G**) in (9) is relevant only when **G** is singular.

The original approach of Hansen & Voje (2011) was to calculate *e*(**x**) or *c*(**x**) along the observed change and compare it with its maximum, minimum, and mean under the uniform distribution of **x** as described by Hansen & Houle (2008). Hunt (2007) used Monte Carlo simulations to draw a null distribution of *e*(**x**) under the same condition. These can be replaced by the analytic probability distributions of these quantities as ratios of quadratic forms, assuming the multivariate normality of **x**, for greater accuracy and enhanced power. Moments of *e* and *c* in this general condition can be obtained as described in Watanabe (2023a).

In this procedure, *e*(**x**) should be regarded as a descriptive measure of the amounts of genetic variance in the direction of **x** rather than that originally available to selection. It is possible to insert 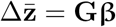 into **x** in order to model the distribution of *e*(**x**) under the Lande model with various selection regimes, as done for *Q*_**V**_ above. On the other hand, interpretation of *c*(**x**) is rather different because it essentially assumes a situation where possible evolutionary change is confined to the direction of selection by strong stabilizing selection in all other directions (Hansen & Houle, 2008; Hansen *et al*., 2019). When strictly applied, therefore, *c*(**x**) is supposed to represent the hypothetical genetic variance that was available to selection. For modeling the distribution of *c*(**x**) under various selection regimes, one needs to insert **β** rather than **Gβ** into **x** in (9).

Example distributions of *e* and *c* with uniformly distributed **x** are shown in Fig. 4, in a setting where the eigenvalues of **G** are geometrically decreasing with *V*_rel_ = 0.2 (as in Fig. 3c). Also shown there is the distribution of *e* along 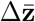 under the Lande model with uniformly distributed **β**; the corresponding distribution for *c* is identical to that with uniformly distributed **x** for the above reason. Examples under different eigenvalue conformations are shown in Supplementary Material (Figs. S1–S4). The eigenvalues are standardized so that the mean *ē* equals 1 across varying *p*. When *p* = 2, the distribution of *e* under the uniform distribution of **x** is a scaled beta distribution (Appendix A). The distributions can be multimodal, as previously mentioned by Watanabe (2023a) as a possibility. These multimodal cases are where inspection of the density, rather than just the mean and range, is most relevant.

**Figure 4.**
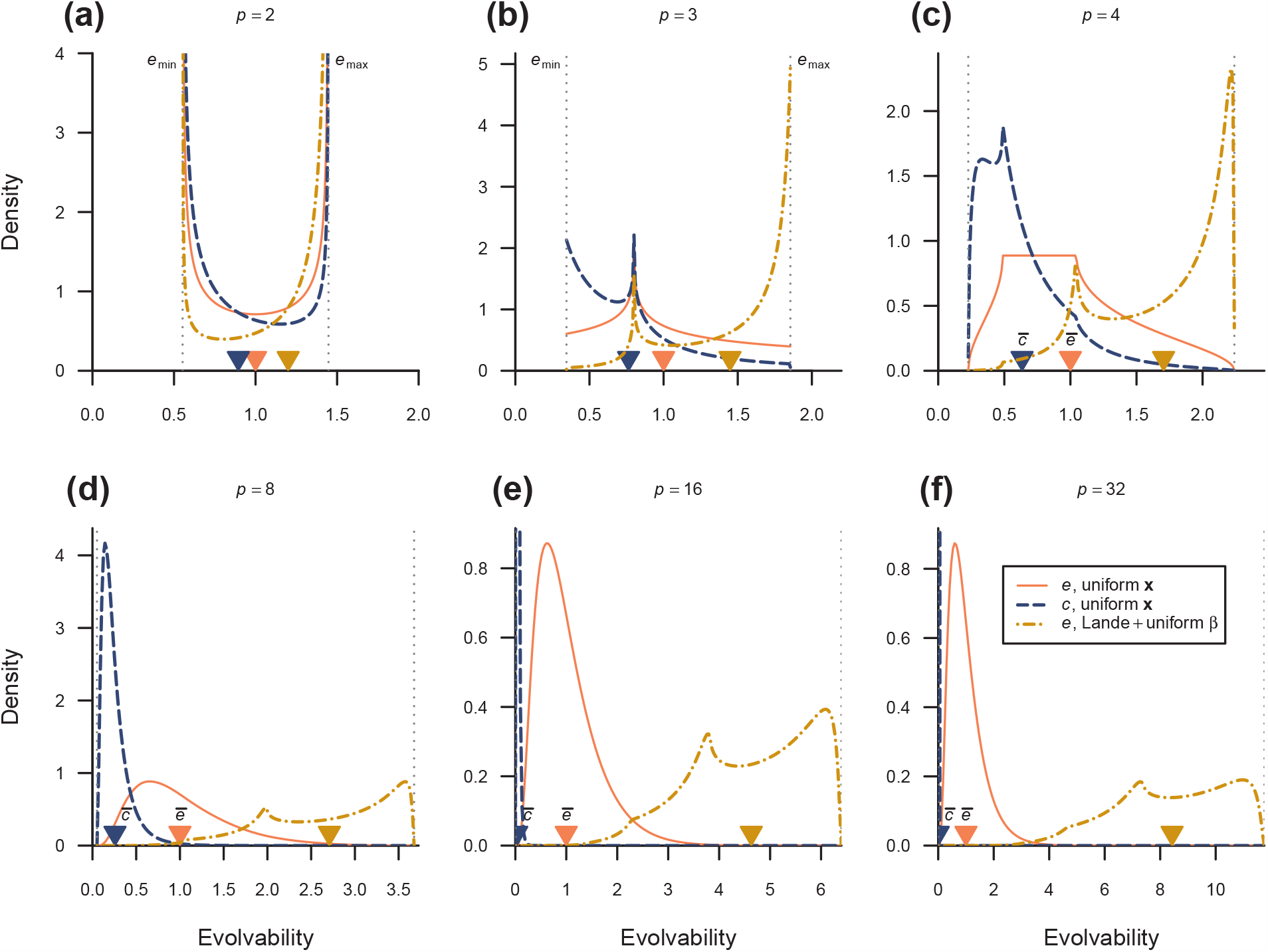
Example distributions of evolvability. Densities of evolvability *e* (orange solid line) and conditional evolvability *c* (blue broken line) for uniformly distributed vectors, as well as that of *e* under the Lande model with uniformly distributed **β** (gold broken line), are shown for the same eigenvalue conformations as in Fig. 3c. Triangles at the bottom represent the means. The density for *c* in **e** and **f** are too strongly concentrated around the mean to be visualized in the same panels. Dotted vertical lines denote the limits of *e* and *c*. Theory states that the density is undefined at the eigenvalues of **G** (Appendix A).

## 3 Empirical examples

Here, distribution theories described above are applied to two previously published datasets, which were chosen to demonstrate how interpretation of results can change by incorporating distribution theories. R scripts to reproduce these reanalyses are available in Supplementary Material.

### 3.1 *Asellus* data

Eroukhmanoff & Svensson (2011) studied phenotypic divergence in two independent invasion from the reed into stonewort habitats by an aquatic isopod *Asellus aquaticus* using four morphometric and three pigmentation-related traits (*p* = 7).^3^ In two ancestor–descendant population pairs (Lake Krankesjön and Lake Tåkern), they estimated **G** from common garden experiments using the animal model, and calculated the angles of deviation *θ* of observed changes in mean phenotypes from the respective **g**_max_. They interpreted the observed angles of 1.12 and 1.29 (in radian; or 64° and 74° as originally presented) in the 7-dimensional space as “almost orthogonal” (p. 1370) and conjectured that the adaptation in this system might not be severely constrained by genetic covariation. From the present distribution theories, however, the squared cosines of these angles (0.192 and 0.076) are not much different from the expectation from the null distribution of uniformity in a 7-dimensional space (1*/*7 *≈* 0.143), with the corresponding lower *P* -values from the beta distribution being 0.723 and 0.491, respectively (Fig. 5a, b). Hence, these deviations are just as expected in a 7-dimensional space and not to be considered particularly large.

**Figure 5.**
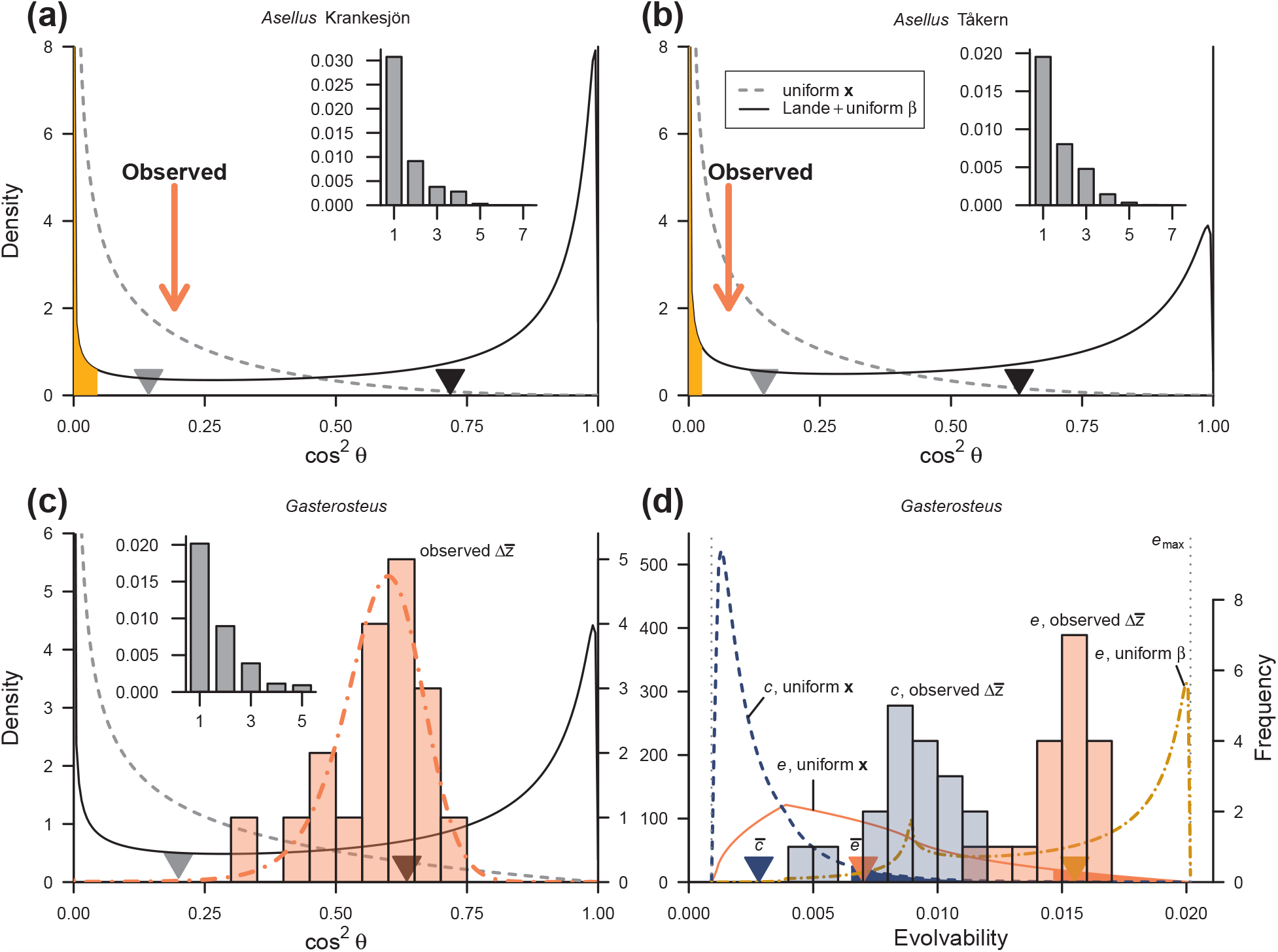
Results of reanalysis. (**a, b**) Squared cosine of angles of deviation from **g**_max_ in two population pairs of *Asellus* from Eroukhmanoff & Svensson (2011). Observed values are indicated in orange arrows and compared with the null distributions (gray broken lines) and the distributions under Lande model assuming uniform **β** (black solid lines), with triangles at the bottom denoting means of these distributions. Solid fill on the lower tail denote regions below 5 percentiles for the latter distributions. Inset scree plots show eigenvalue conformations of the respective **G**. (**c**) Same in 18 populations of *Gasterosteus* from Berner *et al*. (2010). Observed values are indicated in density-scaled histograms with the corresponding scale on the right-hand side margin. Theoretical distribution of *Q*_**g**_max using the sample mean and covariance of 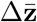 is shown with orange broken line. (**d**) Evolvability in the *Gasterosteus* data. Distributions of evolvability *e* and conditional evolvability *c* under uniform distribution of **x** are shown in orange solid and blue broken lines, respectively, and that of *e* under Lande model assuming uniform **β** is in gold broken line. Solid fills on the upper tails are regions above 95 percentiles.

For rejecting the hypothesis of genetic constraints in the sense of the Lande model, one would look at the position of the squared cosine in the corresponding distribution under the model. In the absence of any a priori hypothesis on the distribution of selection gradients, it seems natural to assume the uniform distribution of **β**. The distribution of squared cosine was drawn for each population pair using the respective **G**, and the resulting lower *P* -values were 0.114 and 0.097 for the Krankesjön and Tåkern, respectively (Fig. 5a, b). This result does not seem to suggest strong evidence against the Lande model, unlike the conjecture of Eroukhmanoff & Svensson (2011). All these said, looking at the angle alone will not be the best way for testing a constraint hypothesis (see Discussion).

### 3.2 *Gasterosteus* data

Berner *et al*. (2010) studied phenotypic divergence in 18 independent invasions from the marine to lacustrine habitats by a stickleback fish *Gasterosteus aculeatus* using four morphometric and one meristic traits (*p* = 5). They used the phenotypic covariance matrix **P** in a marine population as a substitute of ancestral **G** and calculated the angles of deviation *θ* of observed phenotypic changes from its major axis. The observed angles ranged 0.564–0.967 with mean 0.706. Using a bootstrap-based test of Berner (2009), they rejected the hypothesis of collinearity between the divergence and major axis, and concluded that the ancestral trait covariance has not appreciably biased evolutionary trajectories (see Discussion for comments on this test). This procedure was subsequently criticized by Hansen & Voje (2011) who proposed using evolvability measures instead of the angle of deviation (above), but for demonstrative purposes the angle data are re-analyzed first. The squared cosines corresponding to these observations ranged 0.323–0.715 (mean 0.579), all of which are larger than the expectation from random alignment in a 5-dimensional space, 0.200 (Fig. 5c). These values are in fact distributed around the mean 0.635 under the Lande model assuming uniform distribution of **β**. Evidently, however, that particular theoretical distribution does not fit the data well (Kolmogorov–Smirnov test; *D* = 0.534, *P <* 0.001), so an alternative explanation might be sought.

It might be tempting to insert the mean and covariance of the selection gradients **β** reconstructed from observed divergence vectors using the Lande equation 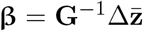. When this is done from the 18 observed divergence vectors and the reconstructed mean and covariance are plugged in as parameters of the multivariate normal distribution, the resultant distribution of *Q*_**g**_max showed an almost perfect fit to the observed data (Kolmogorov–Smirnov test; *D* = 0.108, *P* = 0.970; Fig. 5c). However, this is no more than just using the sample mean and covariance of 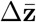 as parameters of **x** in the original definition of *Q*_**g**_max (3) because **G**^*−*1^ cancels with **G** in (6). In other words, this plugged-in distribution does not bear information regarding the fit of the Lande model, so cannot be used to make inference on the biological processes of interest. Nevertheless, it seems notable that the present distribution theory on *Q*_**V**_ assuming multivariate normality of **β** can fit empirical data fairly well in some situations. Thus, the present theory might be of potential use when external estimates are available for the parameters.

Next, following Hansen & Voje (2011), the distributions of evolvability *e* and conditional evolvability *c* in the directions of observed divergences were calculated and compared with their null distributions. As found by Hansen & Voje (2011) using the same procedure with partly different expressions, all observed values (0.0115–0.0164 and 0.0043–0.0119 for *e* and *c*, respectively) were found above the respective expectation *ē* = 0.0070 and 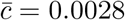 (Fig. 5d). The present theory further indicated that most observed values (13 and 16 out of 18 for *e* and *c*, respectively) have upper *P* -values smaller than 0.05.^4^ This result strengthens the argument of Hansen & Voje (2011) that the evolutionary divergences in this system have likely occurred in directions of rather high evolvability, in contrast to the original argument by Berner *et al*. (2010).

Nevertheless, it is notable that the observed distribution of *e* does not conform with the theoretical distribution of *e* under the Lande model and uniform distribution of **β** (Kolmogorov– Smirnov test; *D* = 0.540, *P <* 0.001). In particular, the observed *e* does not involve any large value close to *e*_max_ which is expected to happen with fair probability under that distribution. It seems reasonable to conclude that the evolution has happened in directions with relatively high, but not the highest, evolvability.

## 4 Discussion

### 4.1 Consideration on sampling error

A major limitation of the above distribution theory is that it has not incorporated potential effects of sampling error in **G** or divergence vectors, practically treating them as fixed. Comprehensive treatment of the effect of errors is beyond the scope of the present study, but several qualitative remarks seem pertinent. First, in most sampling designs, empirical estimates of **G** (or its substitutes; hereafter **Ĝ**) and evolutionary trajectory vectors can likely be considered independent. This is because, at least under multivariate normality of the sample, the sample mean vector and covariance matrix are independent of each other (e.g., Anderson, 2003, chapter 3). As long as the sampling distribution of **Ĝ** is approximated by that of the ordinary sample covariance matrix, this will ensure independence between **Ĝ** and evolutionary trajectory vectors, the latter of which are typically calculated from sample mean vectors. In this case, the effects of sampling can in principle be treated separately between (components of) **Ĝ** and evolutionary trajectory vectors.

If the sampling error in the evolutionary trajectory is multivariate normal, it can in principle be incorporated into the present distribution theory as the parameters in the distribution of **x**. If this sampling error is spherical (i.e., unbiased, independent, and of the same magnitude across traits), it will not affect any results assuming the spherical distribution (or the uniform distribution on *S*^*p−*1^, when scaled) of **x**, effectively leaving the sampling error in **Ĝ** as the only concern. In fact, the null distribution for *Q*_**V**_ (5) is unaffected by that sampling error, because the distribution is a function of *p* and *q* only.

The effects of sampling on the null distributions of *e* or *c* are more complicated, because these are influenced by all eigenvalues of **G**. It is widely recognized that sampling error causes estimation bias in the eigenvalues of a covariance matrix, so that sample eigenvalues are typically, though not always, more widely dispersed than the corresponding parameters (see Marroig *et al*., 2012; Grabowski & Porto, 2017; Watanabe, 2022b, and references therein). If an unbiased estimate **Ĝ** of **G** is available, as for the ordinary covariance matrix, tr **Ĝ** */p* is an unbiased estimate of the average evolvability *ē* = tr **G***/p*. The sampling error in eigenvectors should be irrelevant, as long as it is independent of that of eigenvalues and the distribution of **x** is spherical.

The effect of errors on non-null distributions of *Q*_**V**_ and evolvability measures will be much more complicated. Errors in **x** or eigenvectors of **Ĝ** will usually cause downward bias in *Q*_**V**_ unless *q* is large relative to *p*, because the denominator is more influenced by the error added as sum of squares than the numerator is. Note that this effect grows as *p* increases. This might be counteracted by the upward bias in large eigenvalues of **Ĝ**, so it is rather difficult to make a qualitative surmise on the effect of errors under non-null conditions at present.

### 4.2 Testing constraint hypothesis

For testing the hypothesis of alignment between **g**_max_ and evolutionary trajectory vectors, Schluter (1996; Schluter in Bégin & Roff, 2003) proposed a bootstrap-based test which concerns the null hypothesis that *θ* = 0 and that any observed deviation arises from sampling error. Although such a resampling-based approach will be a promising way to handle sampling error, there is a complexity that the angle cannot be negative, so that any sampling error will cause upward bias in observed angles when *θ* = 0 in the population. That issue was partly addressed by the procedure of Berner (2009) which attempts to correct the bias by subtracting the observed angle from the bootstrap sample. Nevertheless, there remains a fundamental question whether these test procedures address a meaningful hypothesis. Under the Lande model, perfect alignment between **g**_max_ and response (*θ* = 0) is expected only when **g**_max_ and **β** are perfectly aligned. Test for that hypothesis might be of potential interest in certain situations, but that should not be taken as test for a genetic constraint hypothesis. Confusion on this point seems to be the primary source of contrasting interpretations between, e.g., Berner *et al*. (2010) and subsequent reanalyses (Hansen & Voje, 2011; this study), apart from the technical issues mentioned above. Although a large deviation of an evolutionary trajectory from **g**_max_ is sometimes used as evidence against the power of genetic constraints (Berner *et al*., 2010; Eroukhmanoff & Svensson, 2011), examining the angle alone is usually not the best way for testing that hypothesis. A relatively minor reason is that there can be more than one axes with large genetic variance, which can in principle be mitigated (above). A more fundamental reason is that the angle does not bear information on the amount of evolutionary change. This caveat even applies to the evolvability-based approach; evolvability measures as used therein are just descriptive measures of the amount of genetic variance available in the directions of evolutionary trajectories. Obviously, a large deviation from **g**_max_ or small evolvability associated with a relatively small amount of change will still be consistent with a constraint hypothesis. Therefore, test of a genetic constraint hypothesis will need to incorporate the amount of evolutionary change and/or time since divergence, as originally proposed (Schluter, 1996; Hunt, 2007; Hansen & Voje, 2011). In this sense, joint examination of evolvability and the amount of phenotypic divergence (Bolstad *et al*., 2014; Opedal *et al*., 2022) seems to be a sensible approach. Univariate tests based on the present theories will be one of the possible toolkits for testing genetic constraints.

Nevertheless, there can be a practical situation where one must rely on univariate examination of deviation from **g**_max_ or evolvability, e.g., when there is no reliable information on the evolutionary timescale, so that comparison with evolutionary rate is infeasible. In any case, when a univariate test on these measures is required, it is probably worth looking at how clearly the distributions under alternative hypotheses are separated. The present theory clarified that, depending on the eigenvalue conformation of **G**, there can be a large overlap between the relevant distributions (see, e.g., Fig. 4a, b), in which case the univariate test will not be so informative.

### 4.3 Angles in high-dimensional space

The present theories clarify that the distribution of angles depends on the dimensionality of the space *p*. A pair of vectors uniformly distributed in a high-dimensional space tends to be orthogonal to each other as the dimensionality increases. This trend remains true when one of the vectors is replaced by a *q*-dimensional subspace, as long as *p* gets much larger than *q*. Therefore, it is rather misleading to compare angles or vector correlations between datasets with different dimensions. In other words, one should not use intuition in a 2- or 3-dimensional space to interpret these quantities in a high-dimensional space. Some consequences of such practices have been seen in the above empirical examples. Watanabe (2022a) has documented a similar problem in the context of characterizing parallelism between evolutionary trajectories.

This consideration poses a question against the use of angles for measuring deviation between vectors and subspaces. Its use has been encouraged in the biological literature apparently because the angles are supposed to provide intuitive interpretations in high-dimensional geometry that is hard to visualize (e.g., Schluter, 1996; Blows & Walsh, 2009; Bolnick *et al*., 2018). As already seen, intuition from low-dimensional spaces seems to harm more than it benefits. Given that relevant distribution theories are available on (squared) cosines rather than angles, the focus of analysis should probably move from the angles themselves to the cosines or vector correlations, as already practiced by many biologists (Royauté *et al*., 2020; Henry & Stinchcombe, 2023a).

### 4.4 Inference on selection regimes

With the present distribution theory, it becomes possible to evaluate the goodness of fit of observed cosines or evolvability measures to various selection regimes assuming the Lande model and the multivariate normality of the selection gradients. As stated above, however, biologically meaningful inferences cannot be drawn with selection gradients reconstructed from the same observed evolutionary trajectories but require external estimates for the parameters. Therefore, comparison among various selection regimes with this procedure will primarily remain a toolkit for theoretical investigations, unless one has a clear idea on how to obtain statistical parameters from empirical observations.

Ideally, a formal inference on selection regimes should be made without reducing the information into scalar measures like the cosine or evolvability. Inferences based on the entire structure of the covariance matrix of evolutionary trajectories or descendant trait distribution, such as tests for proportionality or identity of covariance matrices (see Ackermann & Cheverud, 2002; Hohenlohe & Arnold, 2008; Revell & Harmon, 2008; Machado, 2020; Mongle *et al*., 2022), would be better suited for that purpose, although those existing methods seem to have their own shortcomings. In this respect, the present procedures using cosines or evolvability measures might remain valid as a heuristic tool to look into evolutionary scenarios underlying phenotypic diversification.

## Acknowledgements

The author would like to thank Masahito Tsuboi for constructive comments. This work was supported by the Newton International Fellowships by the Royal Society (NIF*\*R1*\*180520) and the Overseas Research Fellowships by the Japan Society for the Promotion of Science (202160529).

## APPENDIX A

### Distribution of ratio of quadratic forms

This section provides selected results on the distribution of a (simple) ratio of quadratic forms in normal variables. See, e.g., Mathai & Provost (1992), Paolella (2018, appendix A), and Watanabe (2023b, package vignettes) for more extensive treatments. Let **x** be a *p*-variate random vector, **A** be a *p ×p* symmetric matrix, and **B** be a *p ×p* nonnegative definite matrix. Consider the probability distribution of the following ratio:

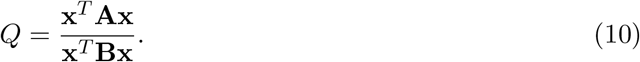

It is well known that, when **B** is nonsingular, *Q* is bounded by the largest *η*_1_ and smallest *η*_*s*_ eigenvalues of **B**^*−*1^**A**: *η*_*s*_ ≤ *Q* ≤ *η*_1_ (e.g., Schott, 2016, section 3.6).

#### A.1 Distribution function

It is seen that the (cumulative) distribution function of *Q* at 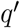 is

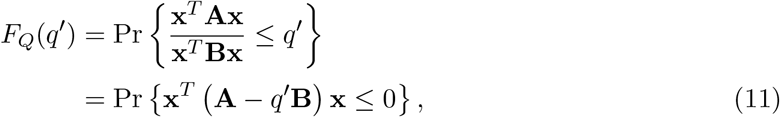

since **x**^*T*^ **Bx** *≥* 0 by assumption. That is, 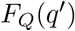 can be expressed by the distribution function of the quadratic form 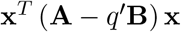 at 0 (see Mathai & Provost, 1992; Forchini, 2002).

Hereafter, assume **x** *∼ N*_*p*_(**µ, I**_*p*_), a *p*-variate normal variable with the mean vector **µ** and the covariance matrix **I**_*p*_.^5^ Forchini (2002, 2005) derived explicit expressions of 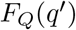 in this condition. Let **H** be a matrix of eigenvectors of 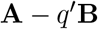 such that

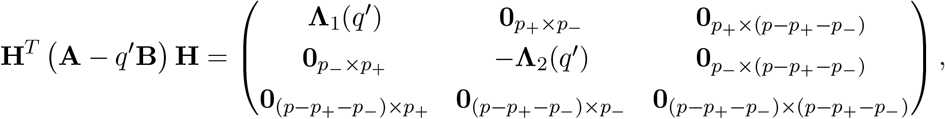

where **Λ**_1_(*q*^*′*^) and *−***Λ**_2_(*q*^*′*^) are *p*_+_- and *p*_*−*_-dimensional diagonal matrices of positive and negative eigenvalues of 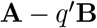, respectively, and **0**_*·×·*_ denote matrices of zeros of appropriate dimensions (which can disappear when *p*_+_ + *p*_*−*_ = *p*). Note that, from (11), 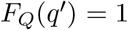 when 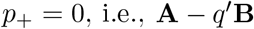 is negative (semi)definite; similarly, 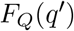 = 0 when *p*_*−*_ = 0. This situation is when 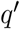 is outside the bound of eigenvalues mentioned above. By denoting

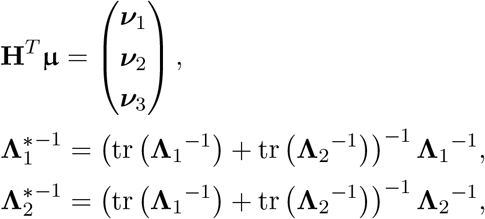

with tr (*·*) being matrix trace and the partition of **H**^*T*^ **µ** corresponding to that of the rows of 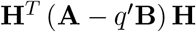 above, the expression of Forchini (2005, after correcting some errors) is

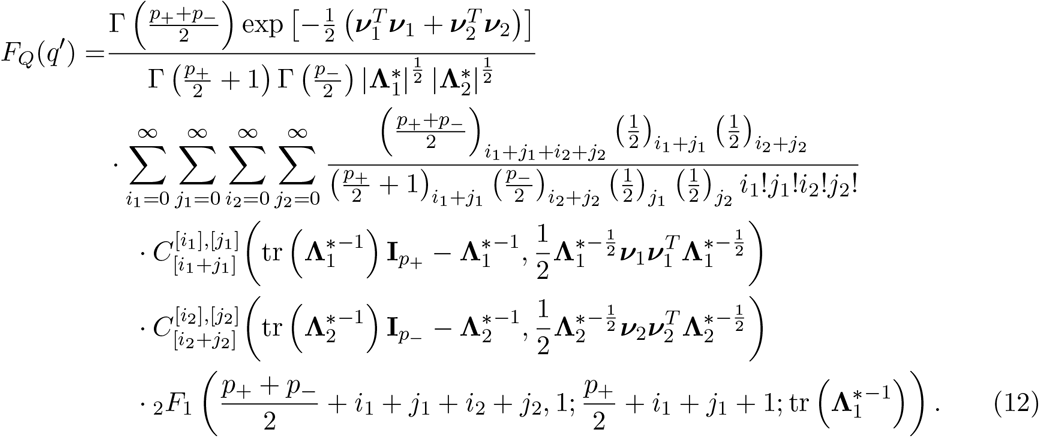

Here, Γ (*·*) is the gamma function, (*·*)*k* denotes Pochhammer’s symbol, (*a*)*k* = *a*(*a*+1) … (*a*+*k−*1) (with (*a*)_0_ = 1), |*·*| denotes matrix determinant, and _2_*F*_1_ (*a, b*; *c*; *·*) denotes the (Gauss) hyperge-ometric function. 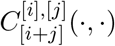 are the top-order invariant polynomials of two matrix arguments of the (*i, j*)-th degree; these are certain scalar-valued functions (homogeneous polynomials) of matrix elements (see Watanabe, 2023a, appendix A for a brief introduction). In the central case (**µ** = **0**_*p*_), the expression simplifies to that of Forchini (2002, theorem 4). The expression of (12) is not particularly enlightening in itself, but can practically be evaluated as a partial sum of the series up to some higher-order terms, with the aid of a recursive algorithm for calculating the top-order invariant polynomials by Hillier *et al*. (2009, 2014).

Alternatively, the distribution function can be evaluated from that of the indefinite quadratic form 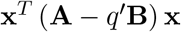 as a linear combination of independent (and potentially noncentral) chi-square variables, by numerical inversion of the characteristic function using the formula of Imhof (1961) (see also Davies, 1973, 1980). Apart from those exact methods, Butler & Paolella (2007, 2008) presented saddlepoint approximations of the distribution function. All these methods have been implemented in the `R` package `qfratio` (Watanabe, 2023b), with the numerical inversion algorithms partly dependent on the package `CompQuadForm` (Duchesne & Micheaux, 2010).

#### A.2 Density function

Explicit forms for the probability density of the ratio of quadratic forms are known only under a restrictive condition. In (10), assume **B** = **I**_*p*_ and **x** is from any spherically symmetric distribution (including, but not limited to, **x** *∼ N*_*p*_(**0**_*p*_, **I**_*p*_)). Under this condition, Hillier (2001) derived an expression for the density, which has different functional forms across intervals bounded by the eigenvalues of **A**. Let *η*_1_, *η*_2_, …, *η*_*s*_ denote the *distinct* eigenvalues of **A** arranged in decreasing order, and *p*_1_, *p*_2_, …, *p*_*s*_ be the corresponding degrees of multiplicity 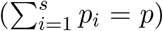 . Consider the transformed variable 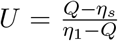 and parameters 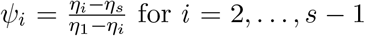, assuming *s >* 2 (see below for the case when *s* = 2).

When *ψ* _*r*+2_ *< u < ψ* _*r*+1_, *r* = 1, …, *s −* 2, the density of *U* at *u* is (from Hillier, 2001, lemma 4)

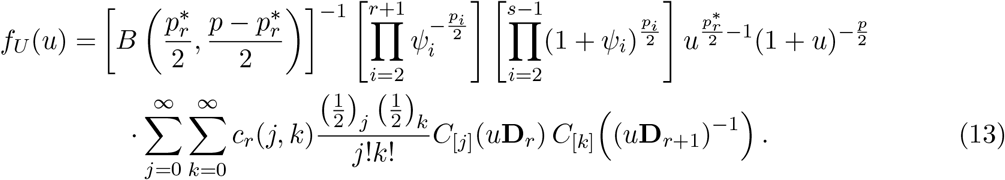

Here, *B* (*·, ·*) is the beta function, 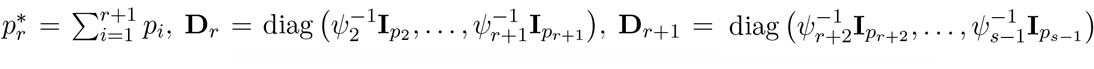, and *c*_*r*_(*j, k*) are the coefficients defined as

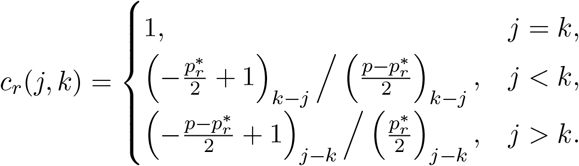

*C*[*j*](*·*) are the top-order zonal polynomials; these are a special case of the invariant polynomials mentioned above with only one argument and are certain homogeneous polynomials of the eigenvalues of the argument matrix.

When 0 *< u < ψ*_*s−*1_ or *ψ*_2_ *< u*, the density is (from Hillier, 2001, lemma 3)

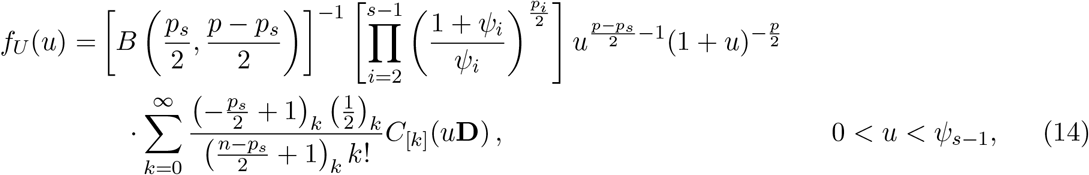

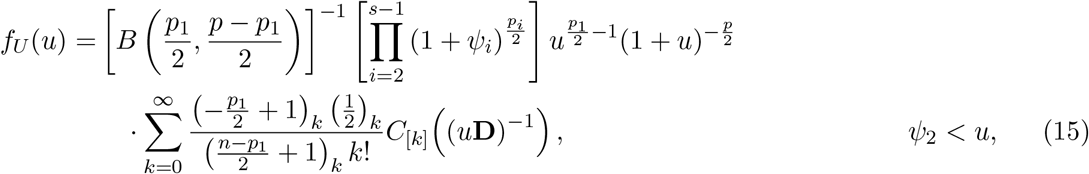

with **D** = diag 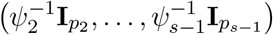. The density is undefined at *u* = *ψ*_*i*_ for any *i*.

From (13), (14), or (15), the density of *Q* at *q*^*′*^ can be obtained as

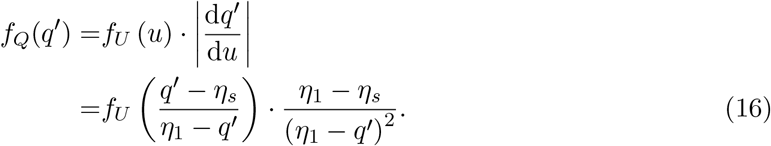

When *s* = 2 (i.e., there are only two distinct eigenvalues), (16) becomes

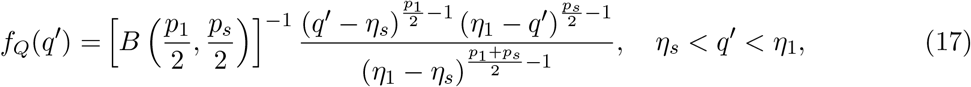

which is the density of the (scaled) beta distribution in the interval (*η*_*s*_, *η*_1_) with the parameters *p*_1_*/*2 and *p*_*s*_*/*2 (see, e.g., Johnson *et al*., 1995). This is in fact expected from the basic relationship between the chi-square and beta distributions mentioned above. Note that (5) is a special case with *η*_1_ = 1 and *η*_*s*_ = 0.

These expressions cannot be used when **B***/*= **I**_*p*_ or **µ***/*= **0**_*p*_. In that case, the only available way to evaluate the exact density seems to be, apart from numerical derivation of the distribution function, conducting numerical inversion using Geary’s formula as documented by Broda & Paolella (2009). Butler & Paolella (2007, 2008) presented saddlepoint approximations of the density. The package `qfratio` implements all these algorithms.

#### A.3 Moments

There is a vast body of literature regarding the moments of a ratio of quadratic forms in normal variables (see Watanabe, 2023a, and references therein). Among several possible expressions, the one with a series expression with the invariant polynomials of matrix arguments seems useful for practical purposes (Hillier *et al*., 2009, 2014; Bao & Kan, 2013). For instance, an expression for the mean when **µ** = **0**_*p*_ is (Smith, 1989; Bao & Kan, 2013)

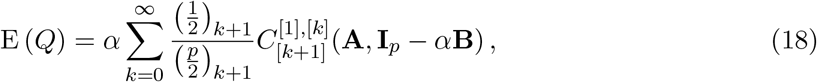

where *α* is any positive constant that is smaller than twice the largest eigenvalue of **B**. When **B** = **I**_*p*_, the series disappears except the first term, and the entire expression simplifies into E (*Q*) = tr **A***/p*, as is well known. See references above for expressions for higher-order moments or the noncentral case **µ***/*≠ **0**_*p*_.

In practice, the series expression can be accurately approximated by its partial sum. The package qfratio (Watanabe, 2023b) implements this method for virtually any possible conditions.

A relevant application is an expression for the mean constraints, which is defined as the mean of vector correlation (cosine) between **g**_max_ and 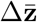 under the uniform distribution of **β** and has been evaluated by Monte Carlo simulations (Marroig *et al*., 2009; Melo *et al*., 2016). The expression is

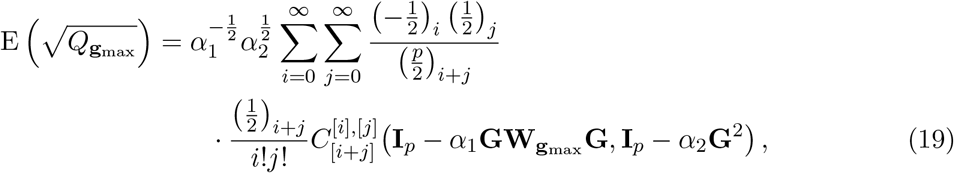

where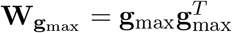, and *α*_1_ and *α*_2_ are any positive constants that satisfy 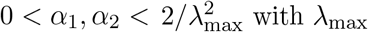 with *λ*_max_ being the largest eigenvalue of **G**. Practically speaking, (19) tends to be rather slow to converge because **I**_*p*_ *− α*_1_**GW**_**g**_max **G** has *p −* 1 eigenvalues of 1. The mean of squared constraints is easier to evaluate, as it involves only a single series. The expression can be obtained by inserting **GW**_**g**_ max **G** and **G**^2^ into **A** and **B**, respectively, in (18). Watanabe (2023a) stated expressions for other mean evolvability measures using the same theory.

## Footnotes

1 Of course, the absolute value of the cosine bears equivalent information, but is not used here because its distribution is (slightly) more cumbersome to characterize.

2 Some biologists have, perhaps unknowingly, drawn vectors from the uniform distribution within a hypercube instead, by using the uniform distribution in [*−*1, 1] as the marginal distributions (McGlothlin *et al*., 2018; Walter, 2023). That distribution does not yield uniformly directed vectors, because there are more vectors directed toward the vertices than toward the faces (see also Rohlf, 2017). It is questionable whether its use can be justifiable from any geometric or biological perspective.

3 It is in general questionable whether covariance between traits measured in different units can be meaningfully interpreted, but this point is ignored for the demonstrative analyses here.

4 Rigorous error rate control for multiple testing will not be necessary for this demonstrative analysis.

5 When **x** *∼ Np*(**µ, Σ**) with a general covariance matrix **Σ**, consider such a *p ×r* matrix that satisfies **KK**^*T*^ = **Σ**, with *r* being the rank of **Σ**, and assume at least one of the following conditions is satisfied: (i) **µ** *∈ R*(**Σ**); (ii) *R*(**A**) *⊆; R*(**Σ**) and *R*(**B**) *⊆; R*(**Σ**); or (iii) **Aµ** = **Bµ** = **0***p*. Then the results can be applied with the following transformed variables and parameters: **x**new = **K**^*−*^**x, µ**_new_ = **K**^*−*^**µ, A**new = **K AK**, and **B**new = **K BK**, where *T T* **K**^*−*^ denotes a generalized inverse of **K** (see Watanabe, 2023a, appendix C). Note that the condition (i) is always satisfied when **µ** = **0***p* or **Σ** is nonsingular, so will cover most cases of practical interest. As stated above, the normality assumption can be released as long as the distribution of **x** is spherically symmetric with E (**x**) = **0***p* because the distribution of *Q* is independent of *∥***x***∥* in this case.

## Notes

### Competing Interest Statement

The authors have declared no competing interest.

