## Supplementary material for "Distribution theories for genetic line of least resistance and evolvability measures": data_B2010.R

```
#####
## This file is part of Supplementary Material for
##   Distribution theories for genetic line of least resistance and
##   evolvability measures
## Junya Watanabe - University of Cambridge, UK.
##
## Gasterosteus data from
## Berner, D., et al. (2010; Evolution)
## https://doi.org/10.1111/j.1558-5646.2010.00982.x

## G matrix of marine (ancestral) population from
## Berner et al. (2010, appendix 1)
G_B2010_m <- matrix(c(
     1.213, -0.110,  0.302,  0.087,  1.356,
    -0.110,  1.012, -0.779, -0.025, -0.358,
     0.302, -0.779, 12.068,  0.170,  5.051,
     0.087, -0.025,  0.170,  3.916, -0.443,
     1.356, -0.358,  5.051, -0.443, 16.840
) / 1000, 5, 5, byrow = TRUE)
rownames(G_B2010_m) <- colnames(G_B2010_m) <- c("TL", "BD", "GW", "RN", "RL")

## Matrix of divergence vectors
## Calculated from ancestral and descendant population means
## From Berner et al. (2010, Supporting Table S2)
delta_zs_B2010 <- matrix(c(
     0.0172,  0.0123,  0.0047,  0.0458, -0.0074,
    -0.0243,  0.0276,  0.0128,  0.0288, -0.1451,
    -0.0079, -0.0020,  0.0886, -0.0716, -0.1800,
     0.0266, -0.0343, -0.0106,  0.0857, -0.0743,
    -0.0136,  0.0025, -0.0228,  0.0375, -0.0944,
    -0.0177, -0.0092,  0.0140,  0.0156, -0.1160,
    -0.0092, -0.0416, -0.0245, -0.0052, -0.0922,
     0.0241,  0.0045, -0.0670, -0.0143,  0.0258,
     0.0354, -0.0258,  0.0025, -0.0179,  0.1862,
     0.0186,  0.0140,  0.0090,  0.0253, -0.0659,
    -0.0098,  0.0139, -0.0204,  0.0024,  0.0891,
     0.0035,  0.0299, -0.0451, -0.0523,  0.0954,
     0.0063, -0.0211,  0.0218,  0.0769, -0.0135,
    -0.0129, -0.0406, -0.0072,  0.0568,  0.0771,
    -0.0120,  0.0010, -0.0179,  0.0504, -0.0412,
    -0.0085, -0.0229,  0.0353, -0.0418,  0.2235,
     0.0079, -0.0248,  0.0569, -0.0540, -0.0482,
    -0.0043,  0.0643, -0.0220, -0.0574, -0.1073
), ncol = 5, byrow = TRUE)
delta_zs_B2010 <- sweep(delta_zs_B2010, 2,
                        c(-0.0398, 0.0488, -0.0410, -0.0178, 0.4451))

## Matrix of divergence vectors, scaled to unit length
## From Berner et al. (2010, Supporting Table S3)
delta_zs_B2010_sc <- matrix(c(
    -0.123,  0.079, -0.099, -0.137, 0.975,
    -0.026,  0.036, -0.090, -0.078, 0.992,
    -0.050,  0.079, -0.201,  0.084, 0.971,
    -0.123,  0.154, -0.073, -0.191, 0.959,
    -0.048,  0.085, -0.033, -0.102, 0.989,
    -0.039,  0.102, -0.097, -0.059, 0.988,
    -0.056,  0.166, -0.030, -0.023, 0.984,
    -0.150,  0.104,  0.061, -0.008, 0.981,
    -0.266,  0.263, -0.154,  0.000, 0.914,
    -0.112,  0.067, -0.096, -0.083, 0.983,
    -0.083,  0.097, -0.057, -0.056, 0.989,
    -0.122,  0.053,  0.011,  0.097, 0.986,
    -0.096,  0.146, -0.131, -0.197, 0.956,
    -0.069,  0.230, -0.087, -0.192, 0.947,
    -0.056,  0.097, -0.047, -0.138, 0.983,
    -0.126,  0.289, -0.307,  0.097, 0.893,
    -0.093,  0.144, -0.191,  0.071, 0.964,
    -0.064, -0.028, -0.034,  0.071, 0.994
), ncol = 5, byrow = TRUE)

## Scale the norms explicitly
delta_zs_B2010_sc <- sweep(delta_zs_B2010_sc, 1,
                           sqrt(diag(tcrossprod(delta_zs_B2010_sc))), "/")
colnames(delta_zs_B2010_sc) <- c("TL", "BD", "GW", "RN", "RL")
rownames(delta_zs_B2010_sc) <-
     c("Big_Mud", "Blackwater", "Cecil", "Ceddar", "Dugout", "Farewell",
       "First", "Gosling", "Gray", "Little_Mud", "Little_Woss", "McCreight",
       "McNair", "Mohun", "Ormond", "Roberts", "Second", "Snow")
```
