## Supplementary material for "Distribution theories for genetic line of least resistance and evolvability measures": data_ES2011.R

```
#####
## This file is part of Supplementary Material for
##   Distribution theories for genetic line of least resistance and
##   evolvability measures
## Junya Watanabe - University of Cambridge, UK.
##
## Asellus data from
## Eroukhmanoff & Svensson (2011; J Evol Biol)
## https://doi.org/10.1111/j.1420-9101.2011.02270.x

## G matrix of Krankesj\"on reed
## Eroukhmanoff & Svensson (2011, Appendix S1)
G_ES2011_K <- matrix(c(
     0.017470,  0.004523,  0.006289,  0.006953, -0.000710,  0.006078, -0.001650,
     0.004523,  0.002752,  0.003704,  0.004171, -0.000310,  0.002726, -0.001970,
     0.006289,  0.003704,  0.005011,  0.005646, -0.000380,  0.003721, -0.002810,
     0.006953,  0.004171,  0.005646,  0.006402, -0.000400,  0.004201, -0.003020,
    -0.000710, -0.000310, -0.000380, -0.000400,  0.000359, -0.000400,  0.000672,
     0.006078,  0.002726,  0.003721,  0.004201, -0.000400,  0.006369, -0.002800,
    -0.001650, -0.001970, -0.002810, -0.003020,  0.000672, -0.002800,  0.008476
), 7, 7, byrow = TRUE)

## G matrix of T\aakern reed
## Eroukhmanoff & Svensson (2011, Appendix S1)
G_ES2011_T <- matrix(c(
     0.011324,  0.003166,  0.004065,  0.004237, -0.001040,   0.003828, -0.003980,
     0.003166,  0.009190,  0.001154,  0.001200, -0.000280,   0.000853, -0.000950,
     0.004065,  0.001154,  0.001491,  0.001554, -0.000310,   0.001112, -0.001190,
     0.004237,  0.001200,  0.001554,  0.001674, -0.000250,   0.001079, -0.001120,
    -0.001040, -0.000280, -0.000310, -0.000250,  0.0013711, -0.001970,  0.000934,
     0.003828,  0.000853,  0.001112,  0.001079, -0.001970,   0.004998, -0.003420,
    -0.003980, -0.000950, -0.001190, -0.001120,  0.000934,  -0.003420,  0.004115
), 7, 7, byrow = TRUE)

rownames(G_ES2011_K) <- colnames(G_ES2011_K) <- 
    rownames(G_ES2011_T) <- colnames(G_ES2011_T) <- 
    c("L", "W1", "W4", "W7", "H", "S", "V")

## Observed angles for Krankesj\"on and T\aakern in the full 7D space
## Eroukhmanoff & Svensson (2011, caption for Fig. 2)
ang_ES2011_K <- 64 / 180 * pi
ang_ES2011_T <- 74 / 180 * pi
```
