## Supplementary material for "Distribution theories for genetic line of least resistance and evolvability measures": README.txt

###############
## Supplementary Material for
##   Distribution theories for genetic line of least resistance and
##   evolvability measures
## Junya Watanabe - University of Cambridge, UK.
##
## This archive contains a supplementary R markdown file for reproducing
## analyses in the above paper.
##
## Contents:
## - README.txt:    This file
## - data_B2010.R:  Data from Berner et al. (2010) used in reanalysis
## - data_ES2011.R: Data from Eroukhmanoff & Svensson (2011) used in reanalysis
## - suppl.Rmd:     Main R markdown source file
## - utils.R:       Definitions of utility functions
##
## Usage:
## You need the following packages installed:
## - 'qfratio':    To run main analysis
## - 'eigvaldisp': To run some codes to reproduce figures
## - 'rmarkdown':  To render R markdown file
## 'eigvaldisp' can be installed from GitHub:
## R> devtools::install_github("watanabe-j/eigvaldisp")
## To render the R markdown file from command line,
## move to this root directory and run
## $  Rscript -e "rmarkdown::render('suppl.Rmd')"
## Alternatively, enter R (GUI) and
## R> setwd("path/to/this/directory")
## R> rmarkdown::render("suppl.Rmd")
## You are free to use knitr or other renderer as you wish.
##
## The scripts were tested with the following dependencies:
## R              v. 4.3.1 (incl. stats, graphics, grDevices)
## eigvaldisp     v. 0.0.0.9405 (available from GitHub)
## qfratio        v. 1.1.0
## MASS           v. 7.3-60
## Rcpp           v. 1.0.11
## RcppEigen      v. 0.3.3.9.3
## rmarkdown      v. 2.24
###############
