## Supplementary material for "Distribution theories for genetic line of least resistance and evolvability measures": suppl.Rmd

````
---
title: "Supplementary material for
        'Distribution theories for genetic line of least resistance and
        evolvability measures'"
author: "Junya Watanabe"
date:   "2023"
output:
  pdf_document:
    fig_width:  6.30
    fig_height: 4.72
papersize: a4
header-includes:
  - \usepackage[labelfont=bf, labelsep=period]{caption}
  - \DeclareCaptionLabelFormat{Sformat}{#1 S#2}
  - \captionsetup[figure]{labelformat=Sformat}
---


This document shows how to reproduce the reanalysis in the main paper using
the `R` environment, and includes some additional figures to supplement
Figure 4 in text.
There is a separate file archive that involves the R markdown source code
of this document and a few additional `R` source files.


# Reanalysis of *Asellus* data

First, load a required package and source definitions of utility functions.
```{r, preparation1}
library(qfratio)
source("utils.R")
```

$\mathbf{G}$ matrices were read from appendix S1, and observed angles were
from the caption for figure 2 of Eroukhmanoff & Svensson (2011).
These are contained in the file `data_ES2011.R`.

```{r, ES2011_1}
## Source data
source("data_ES2011.R")

## G matrix of Krankesjon reed population
G_ES2011_K * 100

## G matrix of Takern reed population
G_ES2011_T * 100

## Number of variables
(p_ES <- nrow(G_ES2011_K))


## Eigenvalues of G
(L_ES2011_K <- eigen(G_ES2011_K, symmetric = TRUE, only.values = TRUE)$values)
(L_ES2011_T <- eigen(G_ES2011_T, symmetric = TRUE, only.values = TRUE)$values)

```


## Analysis of squared cosine
```{r, ES2011_2}
## Get squared cosines
(cossq_ES2011_K <- cos(ang_ES2011_K)^2)
(cossq_ES2011_T <- cos(ang_ES2011_T)^2)
```

Let us obtain (lower) $P$-values corresponding to the observed squared cosines
in the null distribution (eqn.\ (5) in text).
We first calculate the two parameters for the beta distribution, and then use
`stats::pbeta()` to obtain the desired $P$-values.
```{r, ES2011_3}
## Parameters
a_ES <- 1 / 2          # q / 2
b_ES <- (p_ES - 1) / 2 # (p - q) / 2

## P-values
pbeta(c(K = cossq_ES2011_K, T = cossq_ES2011_T), a_ES, b_ES)
## So these observed squared cosines are greater than
## nearly one-half of random alignments in a 7D space!

## Just as a reference, here is how to calculate the mean and variance of
## this beta distribution from the parameters
a_ES / (a_ES + b_ES) # mean (simply 1 / 7 in this case)
a_ES * b_ES / (a_ES + b_ES)^2 / (a_ES + b_ES + 1) # variance
```

Next, consider the distribution under the Lande model assuming uniform
distribution of selection gradients.
$P$-values for ratios of quadratic forms in normal variables can be
calculated with `qfratio:::pqfr()`, and the moments can be
calculated with `qfratio::qfrm()`.

By default, `pqfr()` calculates the distribution function by numerical inversion
using Imhof's (1961) formula.
`qfrm()` calculates a moment as a partial sum of the infinite series;
the argument `m` determines the order of evaluation.
A larger value yields more accurate result at the cost of computation time.
See documentations of the functions and package vignettes for technical details.

```{r, ES2011_4}
## Construct argument matrices;
## proj_Gsq() makes GWG in eqn. (6) with V taken as gmax by default
A_ES2011_K <- proj_Gsq(G_ES2011_K)
B_ES2011_K <- crossprod(G_ES2011_K)
A_ES2011_T <- proj_Gsq(G_ES2011_T)
B_ES2011_T <- crossprod(G_ES2011_T)

## Obtain P-values
(p_cossq_ES2011_K <- pqfr(cossq_ES2011_K, A = A_ES2011_K, B = B_ES2011_K))
(p_cossq_ES2011_T <- pqfr(cossq_ES2011_T, A = A_ES2011_T, B = B_ES2011_T))

## How to calculate the means of the above distributions
(mean_cossq_ES2011_K <- qfrm(A = A_ES2011_K, B = B_ES2011_K, m = 50000))
(mean_cossq_ES2011_T <- qfrm(A = A_ES2011_T, B = B_ES2011_T, m = 50000))

## How to calculate variance
qfrm(A_ES2011_K, B_ES2011_K, 2, m = 50000)$statistic -
    mean_cossq_ES2011_K$statistic^2
```


# Reanalysis of *Gasterosteus* data

Phenotypic covariance matrix $\mathbf{P}$ as a substitute of $\mathbf{G}$ in
the ancestral population, and divergence vectors (both unscaled and scaled) in
the 18 descendant populations were read from the appendix 1 and
supporting tables S2 and S3 in Berner et al. (2010).
These are contained in the file `data_B2010.R`.

```{r, B_prep}
source("data_B2010.R")

## G matrix of marine (ancestral) population
G_B2010_m * 100

## Number of variables
(p_B <- nrow(G_B2010_m))

## Eigenvalues of G
eig_B2010_m <- eigen(G_B2010_m, symmetric = TRUE)
(L_B2010_m <- eig_B2010_m$values)

## gmax
(gmax_B2010_m <- eig_B2010_m$vectors[, 1])
```


## Squared cosine

The analysis proceeds as above, but we would also conduct Kolmogorov--Smirnov
test to see whether the sample conform with the distribution under the Lande
model assuming uniform distribution of selection gradients.
The observed $\Delta \bar{\mathbf{z}}$ scaled to unit length in
the 18 populations are stored in `delta_zs_B2010_sc`, which is contained
in the file `data_B2010.R` sourced above.

```{r, B_sc1}
## Observed squared cosines in 18 populations
cossq_B2010 <- c(delta_zs_B2010_sc %*% gmax_B2010_m)^2
names(cossq_B2010) <- rownames(delta_zs_B2010_sc)
cossq_B2010

a_B2010 <- 1 / 2
b_B2010 <- (p_B - 1) / 2
## Range of upper p-values in the null distribution
range(pbeta(cossq_B2010, a_B2010, b_B2010, lower.tail = FALSE))

## Range of upper p-values in the distribution under Lande model
## assuming uniform distribution of selection gradients
A_B2010 <- proj_Gsq(G_B2010_m)
B_B2010 <- crossprod(G_B2010_m)
range(pqfr(cossq_B2010, A_B2010, B_B2010, lower.tail = FALSE))

## Mean of this distribution; compare with observed values
qfrm(A_B2010, B_B2010, m = 5000)

## Does observed sample conform with this distribution?
ks.test(cossq_B2010, function(q) pqfr(q, A_B2010, B_B2010))
## No
```

As a reference, let us examine the fit of the theoretical distribution for
squared cosine, with the sample mean and covariance plugged-in as parameters
in the distribution of ratio of quadratic forms in normal variables.
This procedure is equivalent to using reconstructed selection gradients using
the Lande equation.
However, remember that this procedure does not speak about the fit of the Lande
model (see text).

The unscaled $\Delta \bar{\mathbf{z}}$ in the 18 populations are stored in
`delta_zs_B2010`, from which sample mean and covariance are calculated first.
```{r, B_sc2}
## Calculate sample mean and covariance of observed delta_zs
mean_delta_z <- colMeans(delta_zs_B2010)
cov_delta_z <- cov(delta_zs_B2010)

## Use them as parameters in the distribution of ratio of quadratic forms
## in normal variables
ks.test(cossq_B2010, function(q) {
            pqfr(q, A = tcrossprod(gmax_B2010_m), B = diag(p_B),
                 mu = mean_delta_z, Sigma = cov_delta_z)
        })
## Great fit, but this does not speak about fit of Lande model (see text)
```


## Evolvability

The analysis of evolvability can be conducted similarly.
We first calculate evolvability $e$ and conditional evolvability $c$
corresponding to the observed $\Delta \bar{\mathbf{z}}$.
These are then compared to their null distribution corresponding to
the uniform distribution of $\Delta \bar{\mathbf{z}}$ using
`qfratio::pqfr()` and `qfratio::qfrm()`.

```{r, B_e1}
## Evolvability
evol_B2010 <- apply(delta_zs_B2010, 1, function(x) {
                        crossprod(x, crossprod(G_B2010_m, x)) / crossprod(x)
                    })
range(evol_B2010)

## Conditional evolvability
G_B2010_m_inv <- chol2inv(chol(G_B2010_m))
cevo_B2010 <- apply(delta_zs_B2010, 1, function(x) {
                        crossprod(x) / crossprod(x, crossprod(G_B2010_m_inv, x))
                    })
range(cevo_B2010)

## Upper p-values
sort(pqfr(evol_B2010, A = G_B2010_m, B = diag(p_B), lower.tail = FALSE))
sort(pqfr(cevo_B2010, A = diag(p_B), B = G_B2010_m_inv, lower.tail = FALSE))
## Many of these are significantly large when compared to those along
## uniformly distributed vectors

## Mean evolvability assuming uniform distribution of response
qfrm(A = G_B2010_m, B = diag(p_B))

## Mean conditional evolvability
qfrm(A = diag(p_B), B = G_B2010_m_inv, m = 1000)
```

As a reference, this is how to calculate the mean of $e$
assuming uniform distribution of selection gradients and the Lande model.
For this purpose, note that
$\Delta \bar{\mathbf{z}} \sim N_p \left( \mathbf{0}_p, \mathbf{G}^2 \right)$
in this case.
```{r, B_e2}
## Mean evolvability assuming uniform distribution of selection + Lande model
qfrm(A = G_B2010_m, B = diag(p_B), Sigma = crossprod(G_B2010_m), m = 5000)
```


# Figures

Scripts for reproducing Figures 3--5 can be found in the R markdown source file,
but are not echoed in this PDF document to save space.

```{r, graph_summary, eval = FALSE, echo = FALSE, warning = FALSE, fig.cap = "Graphical summary of reanalysis. This figure is identical to Figure 5 in text."}
## Vector of quantiles in (0, 1)
quantiles <- seq.int(.Machine$double.eps ^ (1/4),
                     1 - .Machine$double.eps ^ (1/4), length.out = 200)

## Objects that contain densities and means
dist_cossq_ES2011_K <- dist_cossq(G_ES2011_K, quantiles = quantiles, m = 50000)
dist_cossq_ES2011_T <- dist_cossq(G_ES2011_T, quantiles = quantiles, m = 50000)
dist_cossq_B2010 <- dist_cossq(G_B2010_m, quantiles = quantiles, m = 1000)
dist_evol_B2010 <- dist_evol(G_B2010_m)
dist_evol_B2010_rt <- dist_evol(G_B2010_m,
                                Sigma = crossprod(G_B2010_m))

## Density of squared cosine for Gasterosteus data, using sample mean + cov
dseq_rc <- dqfr(quantiles, A = tcrossprod(gmax_B2010_m), B = diag(p_B),
                mu = mean_delta_z, Sigma = cov_delta_z)

### Here are clumsy scripts to reproduce Fig. 5
### You can just jump to the plots...

## Some aesthetic settings
par(mfrow = c(2, 2), mar = c(4.0, 2.4, 2.7, 2.0), oma = c(0, 1.6, 0, 1.7),
    cex = 0.66)
par(fig = c(0/2 + 0.30, 1/2 - 0.08, 1/2 + 0.28, 2/2 - 0.06), new = TRUE,
    mar = c(0, 0, 0, 0), oma = c(0, 0, 0, 0), cex = 0.5)
plot(1, type = "n", axes = FALSE, xlab = "")

## (a) Asellus, Krankesjon
par(fig = c(0/2, 1/2, 1/2, 2/2), new = TRUE,
    mar = c(4.0, 2.4, 2.7, 2.0), oma = c(0, 1.6, 0, 1.7), cex = 0.66)
plot_dseq(quantiles = quantiles, dseq = dbeta(quantiles, a_ES, b_ES),
          mean = 1 / p_ES, plot_new = TRUE, ylim = c(0, 8),
          main = bquote(italic("Asellus")~~"Krankesj\U00f6n"),
          cex.main = 0.8, cex.axis = 0.8, cex_labs = 0.7,
          col = "gray50", lty = "32", y_m = 0.4, lwd = 1.5)
plot_dseq(dist_cossq_ES2011_K, y_m = 0.4)
add_cossq_arrow(ang_ES2011_K, "Observed", adj_text = c(0.5, -0.4),
                cex_text = 1, length = 0.12)
draw_cp(dist_cossq_ES2011_K, col = "orange")
mtext(paste0("(a)"), 3, at = -0.05, line = 0.8, font = 2)
par(fig = c(0/2 + 0.30, 1/2 - 0.08, 1/2 + 0.28, 2/2 - 0.06), new = TRUE,
    mar = c(0, 0, 0, 0), oma = c(0, 0, 0, 0), cex = 0.5)
scree_plot(L_ES2011_K, xlab = "", ylab = "", line_x = -0.5)

## (b) Asellus, Takern
par(fig = c(1/2, 2/2, 1/2, 2/2), new = TRUE,
    mar = c(4.0, 2.4, 2.7, 2.0), oma = c(0, 1.6, 0, 1.7), cex = 0.66)
plot_dseq(quantiles = quantiles, dseq = dbeta(quantiles, a_ES, b_ES),
          mean = 1 / p_ES, plot_new = TRUE, ylim = c(0, 8),
          main = bquote(italic("Asellus")~~"T\U00E5kern"), ylab = "",
          cex.main = 0.8, cex.axis = 0.8, cex_labs = 0.7,
          col = "gray50", lty = "32", y_m = 0.4, lwd = 1.5)
plot_dseq(dist_cossq_ES2011_T, y_m = 0.4)
add_cossq_arrow(ang_ES2011_T, "Observed", adj_text = c(0.2, -0.4),
                cex_text = 1, length = 0.12)
draw_cp(dist_cossq_ES2011_T, col = "orange")
mtext(paste0("(b)"), 3, at = -0.05, line = 0.8, font = 2)
legend("topleft",
       legend = list(bquote("uniform"~bold(x)),
                     bquote("Lande"+"uniform"~bold(beta))),
       col = c("gray50", "black"), lty = c("32", "solid"), lwd = c(1.5, 1),
       cex = 0.7, inset = 0.07, box.lwd = 0.5)
par(fig = c(1/2 + 0.30, 2/2 - 0.08, 1/2 + 0.28, 2/2 - 0.06), new = TRUE,
    mar = c(0, 0, 0, 0), oma = c(0, 0, 0, 0), cex = 0.5)
scree_plot(L_ES2011_T, xlab = "", ylab = "", line_x = -0.5)

## (c) Gasterosteus, squared cosine
par(fig = c(0/2, 1/2, 0/2, 1/2), new = TRUE,
    mar = c(4.0, 2.4, 2.7, 2.0), oma = c(0, 1.6, 0, 1.7), cex = 0.66)
plot_dseq(quantiles = quantiles, dseq = dbeta(quantiles, a_B2010, b_B2010),
          mean = 1 / p_B, ylim = c(0, 6), plot_new = TRUE,
          main = bquote(italic("Gasterosteus")),
          cex.main = 0.8, cex.axis = 0.8, cex_labs = 0.7,
          col = "gray50", lty = "32", y_m = 0.35, lwd = 1.5)
plot_dseq(dist_cossq_B2010, y_m = 0.3)
hist_cosseq_B2010 <- hist(cossq_B2010, freq = FALSE,
                          col = "#FF634770", add = TRUE)
lines(quantiles, dseq_rc, lty = "1242", col = "tomato", lwd = 2)
histd <- with(hist_cosseq_B2010, density / counts)[1]
axis(4, histd * 0:5, labels = 0:5, las = 1, cex.axis = 0.8)

text(0.66, 5.6, bquote("observed"~Delta*bar(z)), adj = c(0, 0), cex = 0.7)
mtext(paste0("(c)"), 3, at = -0.05, line = 0.8, font = 2)
par(fig = c(0/2 + 0.15, 1/2 - 0.25, 0/2 + 0.28, 1/2 - 0.06), new = TRUE,
    mar = c(0, 0, 0, 0), oma = c(0, 0, 0, 0), cex = 0.5)
scree_plot(L_B2010_m, xlab = "", ylab = "", line_x = -0.5)


## (d) Gasterosteus, evolvability
par(fig = c(0.5, 1, 0, 0.5), new = TRUE,
    mar = c(4.0, 2.4, 2.7, 2.0), oma = c(0, 1.6, 0, 1.7), cex = 0.66)
plot_evol(dist_evol_B2010, plot_new = TRUE,
          y_m = max(dist_evol_B2010$dseq_c) / 20,
          main = bquote(italic("Gasterosteus")),
          cex.main = 0.8, cex.axis = 0.8, cex_labs = 0.7, ylab = "",
          adj_m_label_e = c(1.4, -1.2), adj_m_label_c = c(0.5, -1.2),
          show_lims_label_min = FALSE, show_cp = TRUE)
plot_evol(dist_evol_B2010_rt, plot_new = FALSE,
          y_m = max(dist_evol_B2010$dseq_c) / 20,
          adj_m_label_e = c(0.5, -1.2),
          show_lims_label_min = FALSE, show_cp = FALSE,
          show_dseq_c = FALSE, show_mean_c = FALSE, show_m_label = FALSE,
          col_e = "orange3", lty_e = "1141", lwd_e = 1.5)
hist_evol_B2010 <- hist(evol_B2010, freq = FALSE, col = "#FF634770", add = TRUE)
hist_cevo_B2010 <- hist(cevo_B2010, freq = FALSE, col = "#27408B46", add = TRUE)
histd <- with(hist_evol_B2010, density / counts)[1]
axis(4, histd * seq.int(0, 8, 2), labels = seq.int(0, 8, 2),
     las = 1, cex.axis = 0.8)
text(0.016, 400, bquote(italic(e)*","~"observed"~Delta*bar(z)), adj = c(0.5, 0), cex = 0.7)
text(0.009, 290, bquote(italic(c)*","~"observed"~Delta*bar(z)), adj = c(0.2, 0), cex = 0.7)
text(0.005, 160, bquote(italic(e)*","~"uniform"~bold(x)), adj = c(0.4, -0.5), cex = 0.7)
lines(c(0.005, 0.005), c(160, 114))
text(0.003, 290, bquote(italic(c)*","~"uniform"~bold(x)), adj = c(0, -0.2), cex = 0.7)
lines(c(0.003, 0.0024), c(290, 260))
text(0.0196, 340, bquote(italic(e)*","~"uniform"~beta), adj = c(0.9, -0.2), cex = 0.7)
lines(c(0.0196, 0.0199), c(340, 310))
mtext("Frequency", 4, 2, cex = 0.7)
mtext(paste0("(d)"), 3, at = -(max(L_B2010_m) + min(L_B2010_m)) / 20, line = 0.8, font = 2)
```


<!-- # Additional figures -->

Distributions of evolvability measures under various eigenvalue conformations
are provided here to supplement Figure 4 in text.

```{r, preparation2, echo = FALSE}
## For this part, eigvaldisp::GenCov() is used to generate covariance matrices
## with known eigenvalue conformations.  This is available from GitHub.
# devtools::install_github("watanabe-j/eigvaldisp")
library(eigvaldisp)
```

<!-- ## Squared cosine -->

```{r, fig_3, eval = FALSE, echo = FALSE, warning = FALSE, fig.cap = "Distribution of squared cosine. This is identical to Figure 3 in text."}
## Prepare range of p's and graphical parameters
ps <- c(2, 3, 4, 8, 16, 32, 64)
colg <- rainbow(length(ps))
ltys <- c("solid", "44", "32", "2353", "1242", "4323", "24",
          "3433", "5214", "4626")

## Preparation for aesthetic purposes
par(fig = c(0, 1/3, 0.5, 1), mar = c(3.0, 2.4, 2.7, 1.1),
    oma = c(1.4, 1.8, 0, 0), cex = 0.66)
plot(1, type = "n", axes = FALSE, xlab = "")
par(fig = c(1/3 + 0.13, 1/3 + 0.26, 0.75, 0.95), new = TRUE,
    mar = c(0, 0, 0, 0), cex = 0.5)

## Panel a; null distribution
par(fig = c(0, 1/3, 0.5, 1), new = TRUE, mar = c(3.0, 2.4, 2.7, 1.1),
    oma = c(1.4, 1.8, 0, 0), cex = 0.66)
plot_dseq(NULL, ylim = c(0, 4), plot_new = TRUE,
          main = bquote(~"Uniform"), xlab = "", ylab = "",
          cex.main = 0.8, cex.axis = 0.8)
mtext("Density", 2, 2.5, cex = 0.7)
for (i in seq_along(ps)) {
    quantiles <- seq.int(0, 1, length.out = 200)
    plot_dseq(quantiles = quantiles,
              dseq = dbeta(quantiles, 1/2, (ps[i] - 1) / 2),
              mean = 1 / ps[i],
              lty = ltys[i], col = colg[i],
              y_m = par("usr")[4] * 0.04, cex_m = 1.5)
}
legend("topright", legend = ps, col = colg[seq_along(ps)],
       lty = ltys[seq_along(ps)], inset = 0.1,
       title = "p", title.font = 3, cex = 0.7)
mtext("(a)", 3, at = -0.05, line = 0.8, font = 2)
par(fig = c(1/3 + 0.13, 1/3 + 0.26, 0.77, 0.94), new = TRUE,
    mar = c(0, 0, 0, 0), cex = 0.5)

## Panel b; elongated cov matrix (single large eigenvalue)
par(fig = c(1/3, 2/3, 0.5, 1), new = TRUE, mar = c(3.0, 2.4, 2.7, 1.1),
    oma = c(1.4, 1.8, 0, 0), cex = 0.66)
plot_dseq(NULL, ylim = c(0, 8), plot_new = TRUE,
          main = bquote(italic(V)[rel] == 0.2~~"(elongate)"),
          xlab = "", ylab = "", cex.main = 0.8, cex.axis = 0.8)
for (i in seq_along(ps)) {
    G <- GenCov(ps[i], VR = 0.2, shape = "elongate")
    res <- dist_cossq(G, m = 5e6, check_convergence = TRUE, autoscale_args = 10)
    plot_dseq(res, lty = ltys[i], col = colg[i],
              y_m = par("usr")[4] * 0.04, cex_m = 1.5)
}
mtext("(b)", 3, at = -0.05, line = 0.8, font = 2)
par(fig = c(1/3 + 0.13, 1/3 + 0.26, 0.77, 0.94), new = TRUE,
    mar = c(0, 0, 0, 0), cex = 0.5)
scree_plot(diag(GenCov(8, VR = 0.2, shape = "elongate")),
           xlab = "", ylab = "", line_x = -0.5)

## Panel c; gelometrically decreasing eigenvalues
par(fig = c(2/3, 3/3, 0.5, 1), new = TRUE, mar = c(3.0, 2.4, 2.7, 1.1),
    oma = c(1.4, 1.8, 0, 0), cex = 0.66)
plot_dseq(NULL, ylim = c(0, 4), plot_new = TRUE,
          main = bquote(italic(V)[rel] == 0.2~~"(geometric)"),
          xlab = "", ylab = "", cex.main = 0.8, cex.axis = 0.8)
for (i in seq_along(ps)) {
    G <- GenCov(ps[i], VR = 0.2, shape = "exponentially_decreasing")
    res <- dist_cossq(G, m = 5e5, check_convergence = TRUE)
    plot_dseq(res, lty = ltys[i], col = colg[i],
              y_m = par("usr")[4] * 0.04, cex_m = 1.5)
}
mtext("(c)", 3, at = -0.05, line = 0.8, font = 2)
par(fig = c(2/3 + 0.13, 2/3 + 0.26, 0.77, 0.94), new = TRUE,
    mar = c(0, 0, 0, 0), cex = 0.5)
scree_plot(diag(GenCov(8, VR = 0.2, shape = "exponentially_decreasing")),
           xlab = "", ylab = "", line_x = -0.5)

## Panel d; linearly decreasing eigenvalues
par(fig = c(0, 1/3, 0, 0.5), new = TRUE, mar = c(3.0, 2.4, 2.7, 1.1),
    oma = c(1.4, 1.8, 0, 0), cex = 0.66)
plot_dseq(NULL, ylim = c(0, 4), plot_new = TRUE,
          main = bquote("Linearly decreasing"~lambda),
          xlab = "", ylab = "", cex.main = 0.8, cex.axis = 0.8)
mtext("Density", 2, 2.5, cex = 0.7)
mtext(bquote(cos^2~theta), 1, 3, cex = 0.7)
for (i in seq_along(ps)) {
    G <- GenCov(ps[i], shape = "linearly_decreasing")
    res <- dist_cossq(G, m = 5e4, check_convergence = TRUE)
    plot_dseq(res, lty = ltys[i], col = colg[i],
              y_m = par("usr")[4] * 0.04, cex_m = 1.5)
}
mtext("(d)", 3, at = -0.05, line = 0.8, font = 2)
par(fig = c(0 + 0.13, 0 + 0.26, 0.27, 0.44), new = TRUE,
    mar = c(0, 0, 0, 0), cex = 0.5)
scree_plot(diag(GenCov(8, shape = "linearly_decreasing")),
           xlab = "", ylab = "", line_x = -0.5)

## Panel e; quadratically decreasing eigenvalues
par(fig = c(1/3, 2/3, 0, 0.5), new = TRUE, mar = c(3.0, 2.4, 2.7, 1.1),
    oma = c(1.4, 1.8, 0, 0), cex = 0.66)
plot_dseq(NULL, ylim = c(0, 4), plot_new = TRUE,
          main = bquote("Quadratically decreasing"~lambda),
          xlab = "", ylab = "", cex.main = 0.8, cex.axis = 0.8)
mtext(bquote(cos^2~theta), 1, 3, cex = 0.7)
for (i in seq_along(ps)) {
    G <- GenCov(ps[i], shape = "quadratically_decreasing")
    res <- dist_cossq(G, m = 5e4, check_convergence = TRUE)
    plot_dseq(res, lty = ltys[i], col = colg[i],
              y_m = par("usr")[4] * 0.04, cex_m = 1.5)
}
mtext("(e)", 3, at = -0.05, line = 0.8, font = 2)
par(fig = c(1/3 + 0.13, 1/3 + 0.26, 0.27, 0.44), new = TRUE,
    mar = c(0, 0, 0, 0), cex = 0.5)
scree_plot(diag(GenCov(8, shape = "quadratically_decreasing")),
           xlab = "", ylab = "", line_x = -0.5)

## Panel f; two large eigenvalues; deviation from the plane rather than axis
par(fig = c(2/3, 3/3, 0, 0.5), new = TRUE, mar = c(3.0, 2.4, 2.7, 1.1),
    oma = c(1.4, 1.8, 0, 0), cex = 0.66)
plot_dseq(NULL, ylim = c(0, 8), plot_new = TRUE,
          main = bquote(italic(V)[rel] == 0.1~~"(two large"~lambda:~theta~"from PCs 1-2)"),
          xlab = "", ylab = "", cex.main = 0.8, cex.axis = 0.8)
mtext(bquote(cos^2~theta), 1, 3, cex = 0.7)
for (i in seq_along(ps)[-1]) {
    G <- GenCov(ps[i], VR = 0.1, q = 2, shape = "q-large")
    res <- dist_cossq(G, subspace = 1:2, m = 5e6)
    plot_dseq(res, lty = ltys[i], col = colg[i],
              y_m = par("usr")[4] * 0.04, cex_m = 1.5)
}
mtext("(f)", 3, at = -0.05, line = 0.8, font = 2)
par(fig = c(2/3 + 0.13, 2/3 + 0.26, 0.27, 0.44), new = TRUE,
    mar = c(0, 0, 0, 0), cex = 0.5)
scree_plot(diag(GenCov(8, VR = 0.1, q = 2, shape = "q-large")),
           xlab = "", ylab = "", line_x = -0.5)
```


<!-- ## Evolvability -->


<!-- ### Elongate G matrix (one-large eigenvalue) -->

```{r, fig_elongate, echo = FALSE, warning = FALSE, fig.cap = "Distribution of evolvability in single large eigenvalue conditions (Fig.\ 3b)."}
ps <- c(2, 3, 4, 8, 16, 32)
cols <- c("tomato", "royalblue4", "orange3")
ltys <- c("solid", "51", "1141")
lwds <- c(1, 1.5, 1.5)

par(mfrow = c(2, 3), mar = c(3.0, 2.4, 2.7, 1.1), oma = c(1.4, 1.8, 0, 0),
    cex = 0.66)
for (i in seq_along(ps)) {
    G <- GenCov(ps[i], VR = 0.2, shape = "elongate")
    res <- dist_evol(G, m = 5e5, check_convergence = TRUE, autoscale_args = 10)
    res_rt <- dist_evol(G, m = 5e5, check_convergence = TRUE,
                        Sigma = crossprod(G), autoscale_args = 10)
    plot_evol(res, plot_new = TRUE,
              ylim = if (i >= 4) c(0, 1.5) else c(0, 4),
              main = bquote(italic(p)==.(ps[i])), xlab = "", ylab = "",
              cex.main = 0.8, cex.axis = 0.8, lty_c = ltys[2],
              y_m = par("usr")[4] * 0.04, cex_m_e = 1.5, cex_m_c = 1.5,
              show_m_label = (i >= 5),
              adj_m_label_e = c(0.5, -1),
              adj_m_label_c = if (i == 4) c(0.8, -1) 
                              else if (i >= 5) c(1.3, -1) else c(0.5, -1),
              show_lims_label = TRUE,
              adj_lims_label_min = if (i <= 4) c(1.2, 1.2) else c(-0.2, 1.2),
              adj_lims_label_max = if (i <= 4) c(-0.2, 1.2) else c(1.2, 1.2))
    plot_evol(res_rt, plot_new = FALSE,
              cex.main = 0.8, cex.axis = 0.8, lty_e = ltys[3],
              y_m = par("usr")[4] * 0.04, cex_m_e = 1.5,
              show_m_label = FALSE, show_dseq_c = FALSE, show_mean_c = FALSE,
              col_e = cols[3], lwd_e = lwds[3])
    if (i %% 3 == 1) mtext("Density", 2, 2.5, cex = 0.7)
    mtext(paste0("(", letters[i], ")"), 3, at = -0.05, line = 0.8, font = 2)
    if (i >= 4) mtext(bquote(Evolvability), 1, 3, cex = 0.7)
    if (i == 6) legend("topright", inset = 0.12,
                       legend = list(bquote(italic(e)*","~"uniform"~bold(x)),
                                     bquote(italic(c)*","~"uniform"~bold(x)),
                                     bquote(italic(e)*","~"Lande"+"uniform"~bold(beta))),
                       col = cols, lty = ltys, lwd = lwds, cex = 0.7)
}
```

<!-- ### Geometrically decreasing eigenvalues -->
```{r, fig_4, eval = FALSE, echo = FALSE, warning = FALSE, fig.cap = "Distribution of evolvability in geometrically decreasing eigenvalue conditions (Fig.\ 3c). This figure is identical to Figure 4 in text."}
par(mfrow = c(2, 3), mar = c(3.0, 2.4, 2.7, 1.1), oma = c(1.4, 1.8, 0, 0),
    cex = 0.66)
for (i in seq_along(ps)) {
    G <- GenCov(ps[i], VR = 0.2, shape = "exponentially_decreasing")
    res <- dist_evol(G, m = 5e5, check_convergence = TRUE)
    res_rt <- dist_evol(G, m = 5e5, check_convergence = TRUE,
                        Sigma = crossprod(G))
    plot_evol(res, plot_new = TRUE,
              ylim = if (i == 1) c(0, 4)
                     else if (i >= 5) c(0, max(res$dseq_e) * 1.05)
                     else c(0, max(c(res$dseq_e, res$dseq_c, res_rt$dseq_e)) * 1.05),
              main = bquote(italic(p)==.(ps[i])), xlab = "", ylab = "",
              show_m_label = (i >= 3),
              cex.main = 0.8, cex.axis = 0.8, lty_c = ltys[2],
              y_m = par("usr")[4] * 0.04, cex_m_e = 1.5, cex_m_c = 1.5,
              adj_m_label_e = c(0.5, -1),
              adj_m_label_c = if (i >= 4) c(-0.7, -1) else c(0.5, -1),
              show_lims_label = (i <= 2),
              adj_lims_label_min = c(1.2, 1.2),
              adj_lims_label_max = c(-0.2, 1.2))
    plot_evol(res_rt, plot_new = FALSE,
              cex.main = 0.8, cex.axis = 0.8, lty_e = ltys[3],
              y_m = par("usr")[4] * 0.04, cex_m_e = 1.5,
              show_m_label = FALSE, show_dseq_c = FALSE, show_mean_c = FALSE,
              col_e = cols[3], lwd_e = lwds[3])
    if (i %% 3 == 1) mtext("Density", 2, 2.5, cex = 0.7)
    mtext(paste0("(", letters[i], ")"), 3, at = -0.05, line = 0.8, font = 2)
    if (i >= 4) mtext(bquote(Evolvability), 1, 3, cex = 0.7)
    if (i == 6) legend("topright", inset = 0.12,
                       legend = list(bquote(italic(e)*","~"uniform"~bold(x)),
                                     bquote(italic(c)*","~"uniform"~bold(x)),
                                     bquote(italic(e)*","~"Lande"+"uniform"~bold(beta))),
                       col = cols, lty = ltys, lwd = lwds, cex = 0.7)
}
```


<!-- ### Linearly decreasing eigenvalues -->
```{r, fig_lin, echo = FALSE, warning = FALSE, fig.cap = "Distribution of evolvability in linearly decreasing eigenvalue conditions (Fig.\ 3d)."}
par(mfrow = c(2, 3), mar = c(3.0, 2.4, 2.7, 1.1), oma = c(1.4, 1.8, 0, 0),
    cex = 0.66)
for (i in seq_along(ps)) {
    G <- GenCov(ps[i], shape = "linearly_decreasing")
    res <- dist_evol(G, m = 5e5, check_convergence = TRUE)
    res_rt <- dist_evol(G, m = 5e5, check_convergence = TRUE,
                        Sigma = crossprod(G))
    plot_evol(res, plot_new = TRUE,
              ylim = if (i == 1) c(0, 4) else if (i == 6) c(0, 5.5)
                     else c(0, max(c(res$dseq_e, res$dseq_c, res_rt$dseq_e)) * 1.05),
              main = bquote(italic(p)==.(ps[i])), xlab = "", ylab = "",
              cex.main = 0.8, cex.axis = 0.8, lty_c = ltys[2],
              y_m = par("usr")[4] * 0.04, cex_m_e = 1.5, cex_m_c = 1.5,
              adj_m_label_e = if (i == 5) c(1.3, -1) else c(0.5, -1),
              adj_m_label_c = if (i == 5) c(-0.3, -1)
                              else if (i == 1) c(0.8, -1) else c(0.5, -1),
              show_lims_label = (i <= 5),
              adj_lims_label_min = if (i <= 3) c(1.2, 1.2) else c(-0.2, 1.2),
              adj_lims_label_max = if (i <= 3) c(-0.2, 1.2) else c(1.2, 1.2))
    plot_evol(res_rt, plot_new = FALSE,
              cex.main = 0.8, cex.axis = 0.8, lty_e = ltys[3],
              y_m = par("usr")[4] * 0.04, cex_m_e = 1.5,
              show_m_label = FALSE, show_dseq_c = FALSE, show_mean_c = FALSE,
              col_e = cols[3], lwd_e = lwds[3])
    if (i %% 3 == 1) mtext("Density", 2, 2.5, cex = 0.7)
    mtext(paste0("(", letters[i], ")"), 3, at = -0.05, line = 0.8, font = 2)
    if (i >= 4) mtext(bquote(Evolvability), 1, 3, cex = 0.7)
    if (i == 6) legend("topright", inset = 0.04,
                       legend = list(bquote(italic(e)*","~"uniform"~bold(x)),
                                     bquote(italic(c)*","~"uniform"~bold(x)),
                                     bquote(italic(e)*","~"Lande"+"uniform"~bold(beta))),
                       col = cols, lty = ltys, lwd = lwds, cex = 0.7)
}
```


<!-- ### Quadratically decreasing eigenvalues -->
```{r, fig_qua, echo = FALSE, warning = FALSE, fig.cap = "Distribution of evolvability in quadratically decreasing eigenvalue conditions (Fig.\ 3e)."}
par(mfrow = c(2, 3), mar = c(3.0, 2.4, 2.7, 1.1), oma = c(1.4, 1.8, 0, 0),
    cex = 0.66)
for (i in seq_along(ps)) {
    G <- GenCov(ps[i], shape = "quadratically_decreasing")
    res <- dist_evol(G, m = 5e5, check_convergence = TRUE)
    res_rt <- dist_evol(G, m = 5e5, check_convergence = TRUE,
                        Sigma = crossprod(G))
    plot_evol(res, plot_new = TRUE,
              ylim = if (i == 1) c(0, 4) 
                     else c(0, max(c(res$dseq_e, res$dseq_c, res_rt$dseq_e)) * 1.05),
              main = bquote(italic(p)==.(ps[i])), xlab = "", ylab = "",
              cex.main = 0.8, cex.axis = 0.8, lty_c = ltys[2],
              y_m = par("usr")[4] * 0.04, cex_m_e = 1.5, cex_m_c = 1.5,
              show_m_label = (i >= 3),
              adj_m_label_e = if (i == 3) c(1.5, -1) else c(0.5, -1),
              adj_m_label_c = if (i == 1) c(0.8, -1)
                              else if (i == 4) c(1.3, -1) else c(0.5, -1),
              show_lims_label = (i <= 2),
              adj_lims_label_min = if (i <= 1) c(1.2, 1.2) else c(-0.2, 1.2),
              adj_lims_label_max = if (i <= 1) c(-0.2, 1.2) else c(1.2, 1.2))
    plot_evol(res_rt, plot_new = FALSE,
              cex.main = 0.8, cex.axis = 0.8, lty_e = ltys[3],
              y_m = par("usr")[4] * 0.04, cex_m_e = 1.5,
              show_m_label = FALSE, show_dseq_c = FALSE, show_mean_c = FALSE,
              col_e = cols[3], lwd_e = lwds[3])
    if (i %% 3 == 1) mtext("Density", 2, 2.5, cex = 0.7)
    mtext(paste0("(", letters[i], ")"), 3, at = -0.05, line = 0.8, font = 2)
    if (i >= 4) mtext(bquote(Evolvability), 1, 3, cex = 0.7)
    if (i == 6) legend("topright", inset = 0.12,
                       legend = list(bquote(italic(e)*","~"uniform"~bold(x)),
                                     bquote(italic(c)*","~"uniform"~bold(x)),
                                     bquote(italic(e)*","~"Lande"+"uniform"~bold(beta))),
                       col = cols, lty = ltys, lwd = lwds, cex = 0.7)
}
```


<!-- ### Two-large eigenvalues -->
```{r, fig_2large, echo = FALSE, warning = FALSE, fig.cap = "Distribution of evolvability in two-large eigenvalue conditions (Fig.\ 3f)."}
par(mfrow = c(2, 3), mar = c(3.0, 2.4, 2.7, 1.1), oma = c(1.4, 1.8, 0, 0),
    cex = 0.66)
for (i in seq_along(ps)) {
    G <- GenCov(ps[i], VR = 0.1, q = 2, shape = "q-large")
    res <- dist_evol(G, m = 5e5)
    res_rt <- dist_evol(G, m = 5e5, Sigma = crossprod(G), autoscale_args = 50)
    plot_evol(res, plot_new = TRUE,
              dseq_e = if (i == 1) NULL else res$dseq_e,
              dseq_c = if (i == 1) NULL else res$dseq_c,
              ylim = c(0, 4),
              main = bquote(italic(p)==.(ps[i])), xlab = "", ylab = "",
              cex.main = 0.8, cex.axis = 0.8, lty_c = ltys[2],
              y_m = par("usr")[4] * 0.04, cex_m_e = 1.5, cex_m_c = 1.5,
              adj_m_label_e = if (i == 1 || i == 4) c(-0.3, -1)
                              else if (i == 2) c(1.3, -1) else c(0.5, -1),
              adj_m_label_c = if (i == 1 || i == 6) c(1.3, -1) else c(0.5, -1),
              show_lims_label = TRUE,
              adj_lims_label_min = if (i <= 4) c(1.2, 1.2) else c(-0.2, 1.2),
              adj_lims_label_max = if (i <= 4) c(-0.2, 1.2) else c(1.2, 1.2))
    plot_evol(res_rt, plot_new = FALSE,
              cex.main = 0.8, cex.axis = 0.8, lty_e = ltys[3],
              y_m = par("usr")[4] * 0.04, cex_m_e = 1.5,
              show_m_label = FALSE, show_dseq_c = FALSE, show_mean_c = FALSE,
              col_e = cols[3], lwd_e = lwds[3])
    if (i %% 3 == 1) mtext("Density", 2, 2.5, cex = 0.7)
    mtext(paste0("(", letters[i], ")"), 3, at = -0.05, line = 0.8, font = 2)
    if (i >= 4) mtext(bquote(Evolvability), 1, 3, cex = 0.7)
    if (i == 6) legend("topright", inset = 0.12,
                       legend = list(bquote(italic(e)*","~"uniform"~bold(x)),
                                     bquote(italic(c)*","~"uniform"~bold(x)),
                                     bquote(italic(e)*","~"Lande"+"uniform"~bold(beta))),
                       col = cols, lty = ltys, lwd = lwds, cex = 0.7)
}
```
````
