## Supplementary material for "Distribution theories for genetic line of least resistance and evolvability measures": utils.R

```
#####
## This file is part of Supplementary Material for
##   Distribution theories for genetic line of least resistance and
##   evolvability measures
## Junya Watanabe - University of Cambridge, UK.

scree_plot <- function(L,
                       col = "gray60", barw = 0.7,
                       border = NULL, lty = par("lty"), lwd = par("lwd"),
                       xlim = c(1 - barw / 2, p + barw / 2),
                       ylim = c(0, max(L) * 1.05),
                       xlab = "Eigenvalue index", ylab = "Variance",
                       axis_by = 2, line_x = NA, line_y = NA, ...) {
    p <- length(L)
    plot(L, type = "n", axes = FALSE, xlim = xlim, ylim = ylim,
         yaxs = "i", lend = 1, xlab = xlab, ylab = ylab, ...)
    rect(xleft = seq_len(p) - barw / 2, xright = seq_len(p) + barw / 2,
         ybottom = 0, ytop = L, 
         col = col, border = border, lty = lty, lwd = lwd)
    box(bty = "l")
    axis(1, at = seq.int(1, length(L), axis_by), labels = FALSE)
    axis(1, at = seq.int(1, length(L), axis_by), tick = FALSE, line = line_x)
    axis(2, las = 1, labels = FALSE)
    axis(2, las = 1, tick = FALSE, line = line_y)
}

proj_Gsq <- function(G, subspace = 1L,
                     basis = eigen(G, symmetric = TRUE)$vectors[, subspace]) {
    VtVi <- chol2inv(chol(crossprod(basis)))
    # ## Calculate promat (W), which is slow
    # promat <- basis %*% VtVi %*% t(basis)
    # return(G %*% promat %*% G)
    ## This is faster than calculating promat
    GV <- crossprod(G, basis)
    return(tcrossprod(GV, tcrossprod(GV, VtVi)))
    # ## Alternative
    # return(G %*% qr.fitted(qr(basis), G))
}

dist_cossq <- function(G, A = proj_Gsq(G, basis = basis), B = crossprod(G),
                       subspace = 1L,
                       basis = eigen(G, symmetric = TRUE)$vectors[, subspace],
                       from = .Machine$double.eps ^ (1/4),
                       to = 1 - .Machine$double.eps ^ (1/4), length.out = 200,
                       quantiles = seq.int(from, to, length.out = length.out),
                       method_d = c("dqfr", "grad"), autoscale_args = 1,
                       m = 500, mu = rep.int(0, nv), Sigma = diag(nv), ...) {
    method_d <- match.arg(method_d)
    nv <- dim(A)[1L]
    pseq <- qfratio::pqfr(quantiles, A, B, mu = mu, Sigma = Sigma,
                          autoscale_args = autoscale_args)
    if (method_d == "dqfr") {
        dseq <- qfratio::dqfr(quantiles, A, B, mu = mu, Sigma = Sigma,
                              autoscale_args = autoscale_args)
    } else {
        dseq <- numDeriv::grad(
            function(q) qfratio::pqfr(q, A, B, mu = mu, Sigma = Sigma),
            quantiles, side = c(1, rep.int(NA, length(quantiles) - 2), -1))
    }
    mean_cossq <- qfratio::qfrm(A, B, 1, 1, m = m, mu = mu, Sigma = Sigma, ...)
    invisible(list(A = A, B = B, mu = mu, Sigma = Sigma, quantiles = quantiles,
                   pseq = pseq, dseq = dseq, mean = mean_cossq))
}

plot_dseq <- function(obj, quantiles = obj$quantiles, dseq = obj$dseq,
                      mean. = obj$mean$statistic,
                      xlim = c(0, 1), ylim = c(0, 4),
                      xlab = bquote(cos^2~theta), ylab = "Density",
                      main = NULL, plot_new = dev.cur() == 1,
                      lty = 1, col = "black", lwd = 1,
                      y_m = par("usr")[4] * 0.025,
                      pch_m = 25, cex_m = 2,
                    #   lty_m = lty,  lwd_m = lwd * 2,
                      col_m = col, bg_m = col_m,
                      line_x = NA, line_y = NA,
                      line_xlab = 2.5, line_ylab = 2.5,
                      cex_labs = 1, cex_xlab = cex_labs, cex_ylab = cex_labs,
                      cex.axis = 1, ...) {
    if (plot_new) {
        plot(NULL, type = "n", xlim = xlim, ylim = ylim,
             xaxs = "i", yaxs = "i", xlab = "", ylab = "",
             main = main, axes = FALSE, ...)
        box(bty = "u")
        axis(1, at = 0:4 * 0.25, labels = FALSE)
        axis(1, at = 0:4 * 0.25, tick = FALSE,
             line = line_x, cex.axis = cex.axis)
        axis(2, las = 1, labels = FALSE)
        axis(2, las = 1, tick = FALSE, line = line_y, cex.axis = cex.axis)
        mtext(xlab, 1, line_xlab, cex = cex_xlab)
        mtext(ylab, 2, line_ylab, cex = cex_ylab)
    }
    if (!is.null(dseq)) lines(quantiles, dseq, lty = lty, col = col, lwd = lwd)
    # if (!is.null(mean.)) abline(v = mean., lty = lty_m, col = col_m, lwd = lwd_m)
    if (!is.null(mean.)) {
        points(mean., y_m, pch = pch_m, cex = cex_m, col = col_m, bg = bg_m)
    }
}

add_cossq_arrow <- function(angle, text = NULL, y0, y1, f_y0 = 0.6, f_y1 = 0.25,
                            col = "tomato", lwd = 3,
                            font_text = 2, cex_text = 1.5, adj_text = NULL,
                            ...) {
    if (missing(y0) || missing(y1)) {
        u <- par("usr")
        h <- u[4] - u[3]
        if (missing(y0)) y0 <- u[3] + h * f_y0
        if (missing(y1)) y1 <- u[3] + h * f_y1
    }
    cossq <- cos(angle)^2
    arrows(cossq, y0, cossq, y1, col = col, lwd = lwd, ...)
    if (!is.null(text)) {
        text(cossq, y0, text, pos = if(is.null(adj_text)) 3 else NULL,
             font = font_text, cex = cex_text, adj = adj_text)
    }
}

dist_evol <- function(G, length.out = 200,
                      from = min(L) + .Machine$double.eps ^ (1/4) * mean(L),
                      to = max(L) - .Machine$double.eps ^ (1/4) * mean(L),
                      quantiles = seq.int(from, to, length.out = length.out),
                      method_d = c("dqfr", "grad"), autoscale_args = 1,
                      m = 500, mu = rep.int(0, nv), Sigma = diag(nv), ...) {
    method_d <- match.arg(method_d)
    nv <- dim(G)[1L]
    G <- (G + t(G)) / 2
    L <- eigen(G, symmetric = TRUE, only.values = TRUE)$values
    Gi <- chol2inv(chol(G))
    In <- diag(nv)
    pseq_e <- qfratio::pqfr(quantiles, G, In, mu = mu, Sigma = Sigma,
                            autoscale_args = autoscale_args)
    pseq_c <- qfratio::pqfr(quantiles, In, Gi, mu = mu, Sigma = Sigma,
                            autoscale_args = autoscale_args)
    if (method_d == "dqfr") {
        dseq_e <- qfratio::dqfr(quantiles, G, In, mu = mu, Sigma = Sigma,
                                autoscale_args = autoscale_args)
        dseq_c <- qfratio::dqfr(quantiles, In, Gi, mu = mu, Sigma = Sigma,
                                autoscale_args = autoscale_args)
    } else {
        dseq_e <- numDeriv::grad(
            function(q) qfratio::pqfr(q, G, In, mu = mu, Sigma = Sigma),
            quantiles, side = c(1, rep.int(NA, length(quantiles) - 2), -1))
        dseq_c <- numDeriv::grad(
            function(q) qfratio::pqfr(q, In, Gi, mu = mu, Sigma = Sigma),
            quantiles, side = c(1, rep.int(NA, length(quantiles) - 2), -1))
    }
    mean_e <- qfratio::qfrm(G, In, 1, 1, m = m, mu = mu, Sigma = Sigma)
    mean_c <- qfratio::qfrm(In, Gi, 1, 1, m = m, mu = mu, Sigma = Sigma, ...)
    invisible(list(G = G, Gi = Gi, L = L, mu = mu, Sigma = Sigma,
                   quantiles = quantiles,
                   pseq_e = pseq_e, dseq_e = dseq_e, mean_e = mean_e,
                   pseq_c = pseq_c, dseq_c = dseq_c, mean_c = mean_c))
}

plot_evol <- function(obj, quantiles = obj$quantiles,
                      dseq_e = obj$dseq_e, mean_e = obj$mean_e$statistic,
                      dseq_c = obj$dseq_c, mean_c = obj$mean_c$statistic,
                      xlim = c(0, max(obj$L) + min(obj$L)),
                      ylim = c(0, max(c(dseq_e, dseq_c)) * 1.05),
                      xlab = bquote(Evolvability), ylab = "Density",
                      main = NULL, plot_new = dev.cur() == 1,
                      lty_e = "solid", col_e = "tomato", lwd_e = 1,
                      lty_c = "32", col_c = "royalblue4", lwd_c = 1.5,
                      show_dseq_e = !is.null(dseq_e),
                      show_dseq_c = !is.null(dseq_c),
                      show_mean_e = !is.null(mean_e),
                      show_mean_c = !is.null(mean_e),
                      show_m_label = show_mean_e || show_mean_c,
                      y_m = par("usr")[4] * 0.025,
                      pch_m_e = 25, cex_m_e = 2,
                      pch_m_c = 25, cex_m_c = 2,
                      col_m_e = col_e, bg_m_e = col_m_e,
                      col_m_c = col_c, bg_m_c = col_m_c,
                      adj_m_label = c(0.5, -1.8), cex_m_label = 0.8,
                      adj_m_label_e = adj_m_label, adj_m_label_c = adj_m_label,
                      show_lims = plot_new, show_lims_label = show_lims,
                      show_lims_label_min = show_lims_label,
                      show_lims_label_max = show_lims_label,
                      lty_lims = 3, col_lims = "gray40", lwd_lims = 1,
                      cex_lims = cex_m_label,
                      adj_lims_label_min = c(-0.2, 1.2),
                      adj_lims_label_max = c(1.2, 1.2),
                      show_cp = FALSE, alpha = 0.05, lower.tail = FALSE,
                      col_e_cp = col_e, col_c_cp = col_c, border_cp = NA,
                      line_xaxis = NA, line_yaxis = NA,
                      line_xlab = 2.5, line_ylab = 2.5,
                      cex_labs = 1, cex_xlab = cex_labs, cex_ylab = cex_labs,
                      cex.axis = 1, ...) {
    if (plot_new) {
        plot(NULL, type = "n", xlim = xlim, ylim = ylim,
             xaxs = "i", yaxs = "i", xlab = "", ylab = "",
             main = main, axes = FALSE, ...)
        box(bty = "l")
        axis(1, labels = FALSE)
        axis(1, tick = FALSE, line = line_xaxis, cex.axis = cex.axis)
        axis(2, las = 1, labels = FALSE)
        axis(2, las = 1, tick = FALSE, line = line_yaxis, cex.axis = cex.axis)
        mtext(xlab, 1, line_xlab, cex = cex_xlab)
        mtext(ylab, 2, line_ylab, cex = cex_ylab)
    }
    if (show_dseq_e) {
        lines(quantiles, dseq_e, lty = lty_e, col = col_e, lwd = lwd_e)
        if (show_cp) {
            draw_cp(obj, what = "evol", lower.tail = lower.tail, alpha = alpha,
                    col = col_e_cp, border = border_cp)
        }
    }
    if (show_dseq_c) {
        lines(quantiles, dseq_c, lty = lty_c, col = col_c, lwd = lwd_c)
        if (show_cp) {
            draw_cp(obj, what = "cevo", lower.tail = lower.tail, alpha = alpha,
                    col = col_c_cp, border = border_cp)
        }
    }
    if (show_mean_e) {
        points(mean_e, y_m,
               pch = pch_m_e, cex = cex_m_e, col = col_m_e, bg = bg_m_e)
        if (show_m_label) {
            text(mean_e, y_m, bquote(bar(italic(e))),
                adj = adj_m_label_e, cex = cex_m_label)
        }
    }
    if (show_mean_c) {
        points(mean_c, y_m, pch = pch_m_c,
               cex = cex_m_c, col = col_m_c, bg = bg_m_c)
        if (show_m_label) {
            text(mean_c, y_m, bquote(bar(italic(c))),
                adj = adj_m_label_c, cex = cex_m_label)
        }
    }
    if (show_lims) {
        abline(v = c(min(obj$L), max(obj$L)),
               lty = lty_lims, col = col_lims, lwd = lwd_lims)
    }
    if (show_lims_label_min) {
        text(min(obj$L), par("usr")[4], bquote(italic(e)[min]),
             adj = adj_lims_label_min, cex = cex_lims)
    }
    if (show_lims_label_max) {
        text(max(obj$L), par("usr")[4], bquote(italic(e)[max]),
             adj = adj_lims_label_max, cex = cex_lims)
    }
}

draw_cp <- function(obj, what = c("evol", "cevo", "cossq"),
                    alpha = 0.05, add_seq = TRUE, length.out = 21,
                    method_d = c("dqfr", "grad"), lower.tail = TRUE,
                    col = "tomato", border = NA, ...) {
    select_tail <- function(pseq, alpha, lower.tail) {
        if (lower.tail) pseq < alpha else pseq > 1 - alpha
    }
    get_interval <- function(quantiles, ind, lower.tail) {
        if (lower.tail) {
            quantiles[max(which(ind)) + c(0, 1)]
        } else {
            quantiles[min(which(ind)) + c(-1, 0)]
        }
    }
    method_d <- match.arg(method_d)
    if (!is.null(obj$A)) what <- "cossq"
    what <- match.arg(what)
    if (what == "evol") {
        pseq <- obj$pseq_e
        dseq <- obj$dseq_e
        A <- obj$G
        B <- diag(dim(A)[1])
    } else if (what == "cevo") {
        pseq <- obj$pseq_c
        dseq <- obj$dseq_c
        B <- obj$Gi
        A <- diag(dim(B)[1])
    } else {
        pseq <- obj$pseq
        dseq <- obj$dseq
        A <- obj$A
        B <- obj$B
    }
    quantiles <- obj$quantiles
    ind <- select_tail(pseq, alpha, lower.tail)
    int_q <- get_interval(quantiles, ind, lower.tail)
    dseq <- dseq[ind]
    quantiles <- quantiles[ind]
    if (add_seq) {
        mu <- obj$mu
        Sigma <- obj$Sigma
        cp <- stats::optimize(
            function(q) {
                abs(qfratio::pqfr(q, A, B, mu = mu, Sigma = Sigma,
                              lower.tail = lower.tail) - alpha)
                }, int_q)$minimum
        if (lower.tail) {
            from <- cp
            to <- int_q[2]
            new_q <- seq.int(from, to, length.out = length.out)[-1]
        } else {
            from <- int_q[1]
            to <- cp
            new_q <- seq.int(from, to, length.out = length.out)[-length.out]
        }
        if (method_d == "dqfr") {
            new_d <- qfratio::dqfr(new_q, A, B, mu = mu, Sigma = Sigma)
        } else {
            new_d <- numDeriv::grad(
                function(q) qfratio::pqfr(q, A, B, mu = mu, Sigma = Sigma),
                new_q)
        }
        if (lower.tail) {
            dseq <- c(dseq, new_d)
            quantiles <- c(quantiles, new_q)
        } else {
            dseq <- c(new_d, dseq)
            quantiles <- c(new_q, quantiles)
        }
    }
    quant_plot <- c(quantiles, quantiles[seq.int(length(quantiles), 1)])
    dseq_plot <- c(rep.int(0, length(dseq)), dseq[seq.int(length(dseq), 1)])
    polygon(quant_plot, dseq_plot, col = col, border = border, ...)
}
```
